# Pervasive translation of the downstream ORF from bicistronic mRNAs by human cells: impact of the upstream ORF codon usage and splicing

**DOI:** 10.1101/2024.04.26.591296

**Authors:** Philippe Paget-Bailly, Alexandre Helpiquet, Mathilde Decourcelle, Roxane Bories, Ignacio G. Bravo

## Abstract

In this work, we present a straightforward model to study gene expression regulation from virus-like bicistronic mRNAs in human cells, composed by a short *shble* upstream ORF and a reporter *egfp* downstream ORF. We have engineered thirteen synonymous versions of the *shble* ORF to explore a large parameter space in compositional and folding features, as well as differences in propensity to undergo splicing. Our experimental model focuses on the control and modulation of *shble* translation elongation and their effects on downstream translation: i) regarding translation initiation, all constructs share identical 5’UTR, ribosomal binding site and *shble* start codon context; and ii) regarding *egfp* translation, all constructs share identical *shble* stop codon context, spacer between *shble* and *egfp* and the full *egfp* sequence. Our results show first that the human translation machinery can translate the downstream ORF in bicistronic mRNAs, independently of the characteristics of the upstream ORF. Even if GFP protein levels are hundreds of times lower than SHBLE protein levels, our results challenge the textbook interpretation positing that the eukaryotic translation machinery does not handle the downstream ORF in bicistronic transcripts. Second, we show that synonymous recoding of the upstream ORF determines its own translation efficiency, can uncover cryptic splice signals, and largely conditions the probability of translation from the downstream ORF. Our results are consistent with a leaky scanning mechanism at play for downstream ORF translation from bicistronic mRNAs in human cells.

## Introduction

The multitude of cellular phenotypes found in a multicellular organism, referred as cell types, are the result of a multi-layer cascade of molecular mechanisms during gene expression and biological information transfer (Maniatis et Tasic 2002). In multicellular organisms, cells within a tissue display differences in biochemistry and in cellular status that result in alternative transcription and translation, as well as in post-transcriptional and post-translational modifications, overall expanding the set of proteomes, and thus phenotypes, available for expression from a single genome. In the present study, we have focused on the layer for gene expression regulation at the translation level that is allowed by the presence of multiple Open Reading Frames (ORFs) on the same mRNA.

Translation is the most energy-demanding step in gene expression, and thus is highly regulated by multiple *in-cis* acting sequences and *in-trans* acting factors (Lynch et Marinov 2015). Eukaryotic translation begins with scanning of a mRNA from the 5’ 7-meG cap until the recognition of a suitable start codon by the preinitiation complex (Cigan, Feng, et Donahue 1988; Kozak 1989). Because of this directional scanning, the presence of mRNA secondary structures and regulatory sequences between the 5′ 7-meG cap and the ORF start codon greatly affect translation initiation efficiency (Kozak 1986; Hinnebusch, Ivanov, et Sonenberg 2016). The textbook descriptions for genes in eukaryotic cells posit that they are most often arranged in individual promoters and monocistronic mRNAs, compared to the common polycistronic mRNA arrangement in prokaryotic cells. However, mounting evidence suggests that a large fraction of vertebrate mRNAs (in the range of 50%) possess a short regulatory ORF immediately upstream or overlapping the “main” or “canonical” ORF (Chew, Pauli, et Schier 2016; Calvo, Pagliarini, et Mootha 2009; Lin et al. 2019). Such upstream ORFs are proposed to play regulatory roles on the translation of the downstream ORF, either by *in-cis* mechanisms, whereby the presence of the upstream ORF modulates the probability of the ribosome to engage in translation of the downstream ORF, or by *in-trans* mechanisms mediated by the protein product of the upstream ORF (J. Chen et al. 2020).

Compared to the idealized monocistronic mRNAs found in eukaryotes, many RNA and DNA viruses infecting them produce polycistronic mRNAs containing multiple successive ORFs. This arrangement raises the question about viral gene expression regulation, and indeed, viruses have evolved a large repertoire of mechanisms that manipulate the cellular translation machinery to engage into multiple, independent or sequential, translation events from a single mRNA (Walsh et Mohr 2011). Many viruses display for instance internal ribosomal entry sites (IRES) preceding the ORFs present on a single mRNA molecule, thus allowing for ribosomal recruitment and translation of downstream ORFs. Many other viruses, in contrast, tamper with the ribosomal machinery allowing for non-canonical initiation, elongation or termination (Sorokin et al. 2021). The compositional and structural features of the viral mRNA acting *in-cis*, as well as the biochemical context in the infected cell acting *in-trans*, result in a probabilistic, differential ORF translation that influences viral gene expression pattern in a given cell type, determining a characteristic tropism and cell permission pattern and eventually governing viral life cycle.

In this study we have experimentally addressed the question of to what extent the characteristics of an upstream ORF have an impact on the translation of a downstream ORF in human cells. As a toy model for polycistronic viral mRNAs, we have engineered a heterologous expression system expressing bicistronic mRNA in human cells to explore the impact of nucleotide composition, CUPrefs (Codon Usage Preferences) and presence of splicing events in a short upstream ORF, monitoring the translation of a downstream reporter encoding for a fluorescent protein. Using transient transfection on human cells, we have combined transcriptomic, proteomic and cytometry-based fluorescence analyses we provide qualitative and quantitative descriptions about the interplay of mRNA composition, codon recoding and splicing with the cellular machinery, resulting in translation of the downstream ORF in a bicistronic mRNA.

## Results

### Transfection with synonymous versions of a bicistronic construct does not affect heterologous mRNA levels, but synonymous recoding in the upstream ORF can introduce unpredicted splicing events

The thirteen synonymous versions of the *shble* ORF were chemically synthesized and cloned into the pcDNA3.1-C-EGFP expression vector (Fig S1A). Transfection with all constructs results in the expression of a 1602 nt-long mRNA with the same organization: a 161 nt-long 5’UTR; a 414 nt-long *shble* ORF, including a N-terminal AU1 tag and a C-terminal FLAG tag; a 19-nt long gap; a 714 nt-long *egfp* ORF; and a 288 nt-long 3’UTR. The expression vector directs also the expression of a *neo* mRNA under the control of the SV40 promoter, encoding for a NEO protein conferring resistance to the neomycin antibiotic (Fig S1B). Only the *shble* coding sequences differed synonymously between constructs. Versions *shble*#1 to #6 were engineered for a previous study (Picard et al. 2023) to explore the extremes of CUPrefs and nucleotide composition with regards to the average human ones. The design strategy is detailed in Experimental procedures. Versions *shble*#7 to #13 were selected among a pool of a thousand “guided random” *shble* versions to encompass the CUPrefs and mRNA folding energy spectrum found in actual human mRNA sequences. Thus, the 13 *shble* versions display by design compositional variation in G and C frequency in the third codon position (GC3), CpG dinucleotide frequency, TpA dinucleotide frequency, CUPrefs, as well as in mRNA folding energy (Table S1). We have further evaluated the match between the CUPrefs of each version to the average ones in the human genome using COUSIN (Codon Usage Similarity Index), which allows to differentiate and quantify random codon usage, under-and overmatch to a given reference (Bourret, Alizon, et Bravo 2019). Using these five parameters in a principal component analysis (PCA) allowed for sharp discrimination of versions *shble*#1 to #6 and to a lesser extent for versions *shble*#7 to #13 (Fig 1) with the first and second axis capturing 77.3% and 16.3% of the total variance respectively. We have subsequently used the projections of each version on each of these two axes as composite variables to evaluate the impact of codon against the different levels of experimental data we have generated.

**Figure 1:**
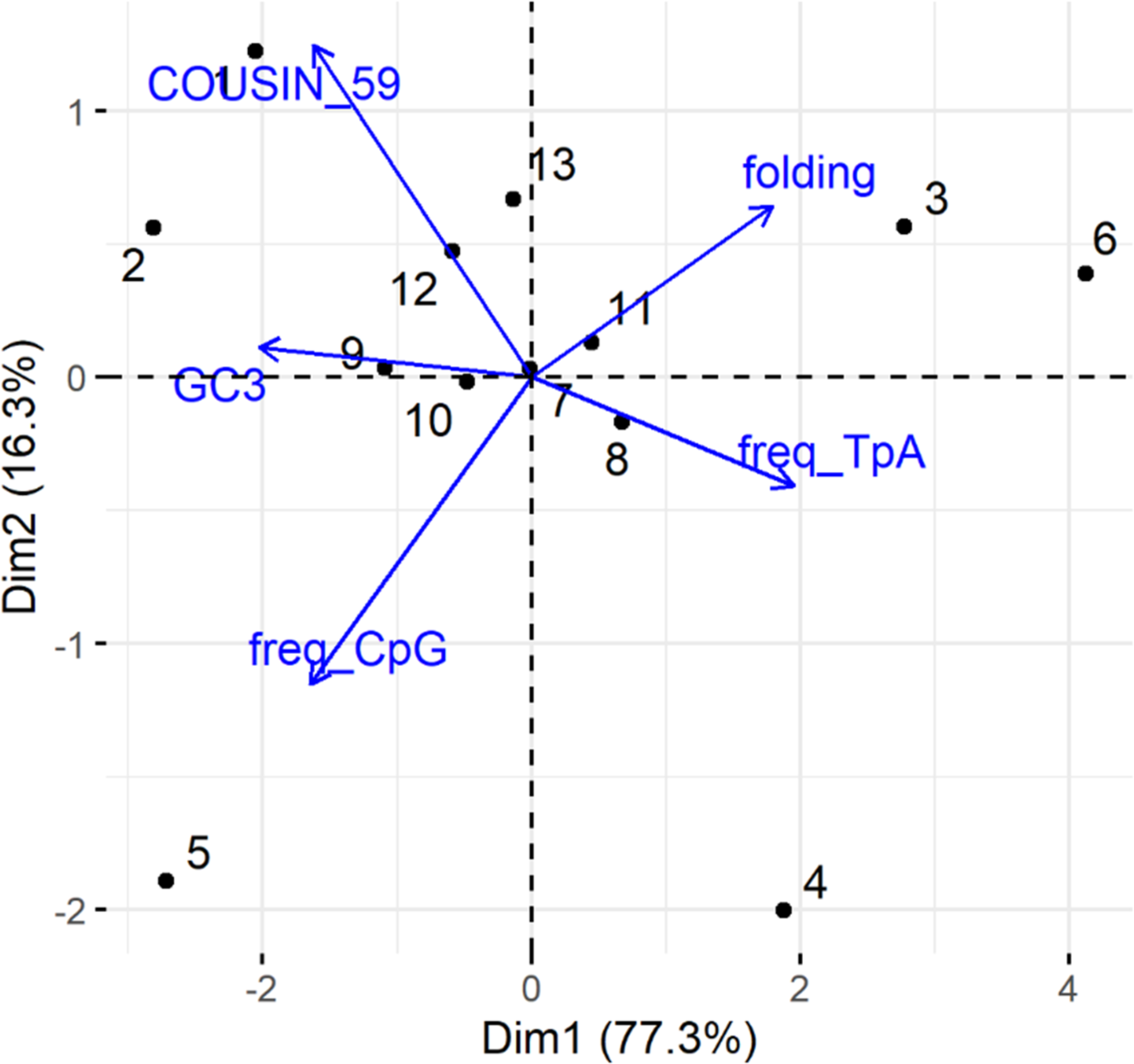
First two axes of a principal component analysis (PCA) of the compositional variables for the thirteen *shble* versions used. The percentage of the total variance captured by each axis is given in parenthesis. GC3, percentage of G or C at the third codon nucleotide codons; freq_CpG, CpG dinucleotide frequency; freq_TpA. TpA dinucleotide frequency; folding, energy of the more stable structure predicted for the mRNA *shble* ORF estimated with UNAfold online tool (http://unafold.org) (Markham et Zuker 2008); COUSIN_59, value of the COdon Usage Similarity Index of the *shble* version with respect to the average human codon usage (Bourret, Alizon, et Bravo 2019).

We identified splicing events in the *shble* ORF when expressing versions *shble*#4, #6 #7, #10 and #13 in the U-2 OS cell line (Fig S2A), even if all thirteen synonymous versions had been scanned for the presence of splice sites using the HSP (Desmet et al. 2009) and SPLM algorithms (Solovyev 2003). Picard and coworkers had previously communicated splicing activity in versions *shble*#4 and #6 when expressed in HEK-293 human cells (Picard et al. 2023). For all spliced versions, we individually identified donor and acceptor sites by RT-PCR followed by Sanger sequencing (Fig. S2B and C). To be able to use these five *shble* versions undergoing splicing to study the impact of mRNA composition and structure on SHBLE and GFP translation, we generated mutated versions of these constructs with ablated splice sites to eliminate splicing as a confounding factor. We took later advantage of the paired spliced/unspliced constructs to study the impact of splicing on translation regulation of our bicistronic mRNAs.

Transient transfection in biological quadru-or octuplicate experiments resulted in normalized mean mRNA levels ranging from 0.80 (95%CI: 0.76-0.84) for *shble*#1 to 1.28 (95%CI: 1.10-1.46) for *shble*#8, as evaluated by RT-qPCR, with no significant differences in mRNA levels among the thirteen constructs (Fig 2A, pairwise Wilcoxon rank sum test with Benjamini-Hochberg (B-H) correction; α=0.05). Only transfection with the control construct, monocistronic *egfp* coding mRNA (mean 1.61, 95%CI: 1.47; 1.74), presented a significant difference with the *shble*#1 condition (Fig 2A, Wilcoxon rank sum test with B-H correction; *p=*0.028). In agreement, variation in mRNA composition and structure did not explain variation in mRNA levels (Fig 2B, R^2^=0, *p=*0.8).

**Figure 2:**
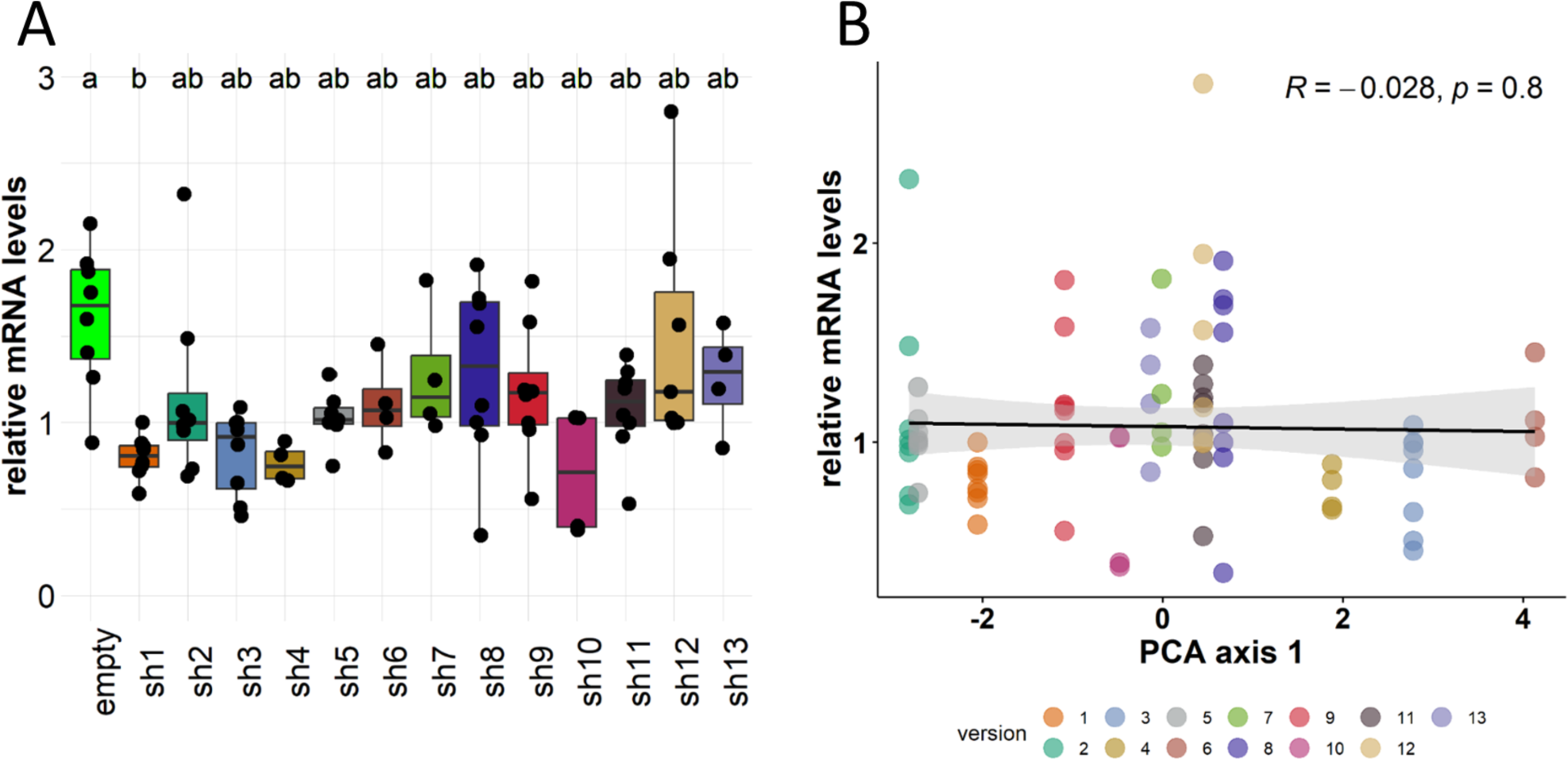
Compositional variation of synonymous *shble* versions does not affect mRNA levels. (panel A) Box-and-whiskers plot showing relative levels of total heterologous mRNAs measured by RT-qPCR from four or eight biological replicates. Within a given replicate, relative mRNA level values were normalised by the median value of all samples. The positive control “empty” condition transcribed a monocistronic *egfp* mRNA while the thirteen *shble* conditions (sh1 to sh13) transcribed a bicistronic *shble_egfp* mRNA. Letters present the results of a pairwise Wilcoxon rank sum test with B-H adjusted p-values. Median values for samples labelled with the same letter are not statistically different (α=0.05). **(panel B)** Pearson’s linear regression (black line) and 95% confidence interval of the fit (grey) between the projection on the first axis of PCA in Fig 1 for each *shble* version, and the relative mRNA levels.

### Human cells constitutively translate the downstream ORF from bicistronic mRNAs, albeit a thousand time less efficiently than the upstream ORF

Label-free proteomic analyses of the thirteen versions in biological triplicates revealed that SHBLE and GFP were respectively on average the 2nd and 1108th most abundant proteins among the 4199 detected, in terms of iBAQ values (the scores correspond to the 8th and 1376th most abundant proteins in raw intensity values; data available in the PRIDE repository entry PXD047576). We verified first that high heterologous protein expression had no impact on the total amount of protein signal detected in each sample (FigS3A) for both SHBLE (FigS3B, R^2^=0, *p=*0.88) and GFP (FigS3C, R^2^=0, *p=*0.93). We then normalised the iBAQ value of SHBLE and GFP proteins by the respective total iBAQ value of each sample to evaluate our target protein levels. We detected SHBLE and GFP protein expression in biological triplicate experiments from all bicistronic mRNAs independently of the *shble* version. SHBLE protein levels were on average 1197 (95%CI: 1131-1364) times higher than GFP levels (Fig. 3, Wilcoxon rank sum test with continuity correction; *p<*3.08*10^-14^). Overall, GFP protein levels produced from the *egfp* monocistronic control were slightly lower than SHBLE protein ones (Fig. 3 Wilcoxon rank sum exact test; *p=*0.030) but most importantly, were significantly higher than GFP protein levels produced from the second ORF of every bicistronic mRNAs (Fig. 3 Wilcoxon rank sum exact test; *p=*1.74*10^-4^). We interpret thus that in bicistronic mRNAs, translation occurs preferentially from the upstream ORF, while translation from the downstream ORF occurs at low levels albeit constitutively.

**Figure 3:**
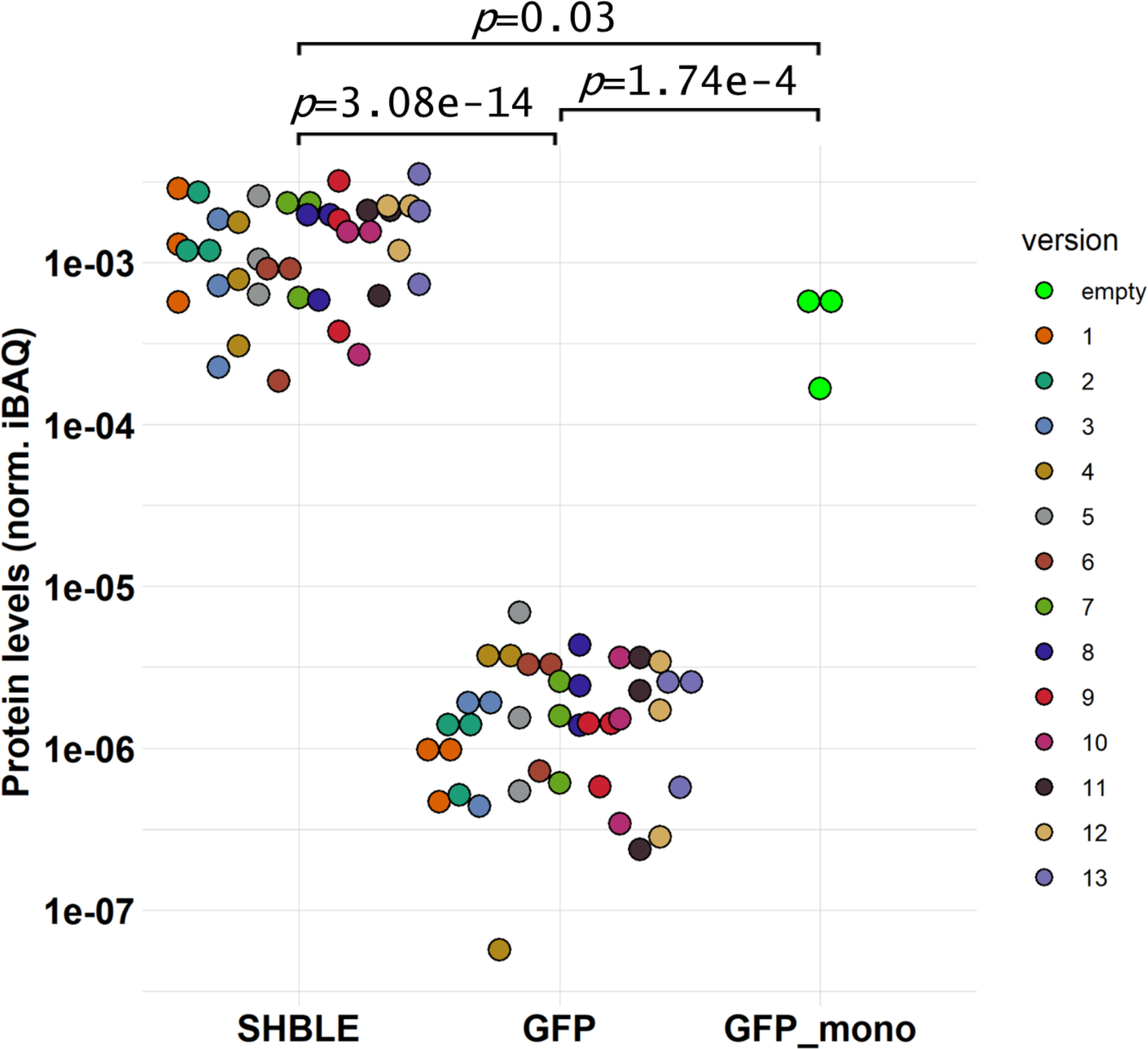
SHBLE and GFP protein levels. Dot-plot showing relative levels of heterologous proteins measured by label-free proteomic from three biological replicates. For each sample, iBAQ values of SHBLE and GFP were normalised by the total iBAQ value of the sample. The positive control “GFP_mono” condition translated GFP from a monocistronic *egfp* mRNA while the thirteen *shble* conditions translated SHBLE and GFP from a bicistronic *shble_egfp* mRNA. GFP values do not follow a normal distribution after a Shapiro normality test (*p*=0.0014), hence the p-values present the results of a pairwise Wilcoxon signed rank exact test.

### Changing nucleotide composition of the upstream *shble* ORF without modifying translation initiation suffices to modulate its own translation efficiency

We quantified next to what extent variation in SHBLE protein levels could be accounted for by variation in mRNA levels and in mRNA composition. Our results using triplicate biological experiments comparing RT-qPCR and proteomic data show that, overall, 71% of the variation in SHBLE levels, can be explained by variation in the bicistronic *shble_egfp* mRNA levels (Fig4A, R^2^=0.71, *p=*1.9*10^-11^).

As a proxy for translation efficiency, we normalised SHBLE protein levels over the corresponding *shble_egfp* mRNA levels. Despite the visual trend for *shble*#1 and #2 to display higher translation efficiency values, and the noteworthy 3-fold variation between *shble*#6 (median 0.40) and *shble*#1 (median 1.30; triplicate median 0.96) (FigS4A), we did not detect significant differences in translation efficiency for SHBLE among the thirteen conditions, most likely because we could use data from only triplicate experiments pairwise (Wilcoxon rank sum test after B-H correction for multiple comparisons; α=0.05). We resorted thus to the analysis of SHBLE translation efficiency as a function of the overall compositional features of the recoded *shble* versions. Our results show that variation in the characteristics of the upstream *shble* ORF explained 44% of the variation in the SHBLE translation efficiency (Fig 4B; R^2^=-0.44, *p=*3.9*10^-6^). The specific analysis by individual variable (Fig S4B to F) showed that SHBLE translation efficiency increased with increased GC3 (R²=0.46, *p=*2.1*10^-6^), with a higher match to the average CUPrefs of the human genome (R²=0.40, *p=*1.7*10^-5^), with increased CpG frequency (R^2^=0.22, *p=*2.5*10^-3^), with decreased TpA levels (R^2^=-0.40, *p=*1.7*10^-5^) and with less stable folding structures of the mRNA *shble* ORF (R^2^=-0.26, *p=*9.9*10^-4^). Finally, we performed a sequential linear regression to determine *a posteriori* which combination of the five parameters used for the PCA in Fig 1 provided the highest explanatory power for variation in SHBLE translation efficiency (Table 1). Sequential linear regression revealed GC3 as the best predictor of SHBLE translation efficiency value (R^2^=0.46), with the four remaining parameters providing no additional explanatory power. When excluding GC3, the best predictor was a combination of TpA frequency, CUPrefs and CpG frequency (R^2^=0.49) with mRNA folding energy not providing additional explanatory power.

**Figure 4:**
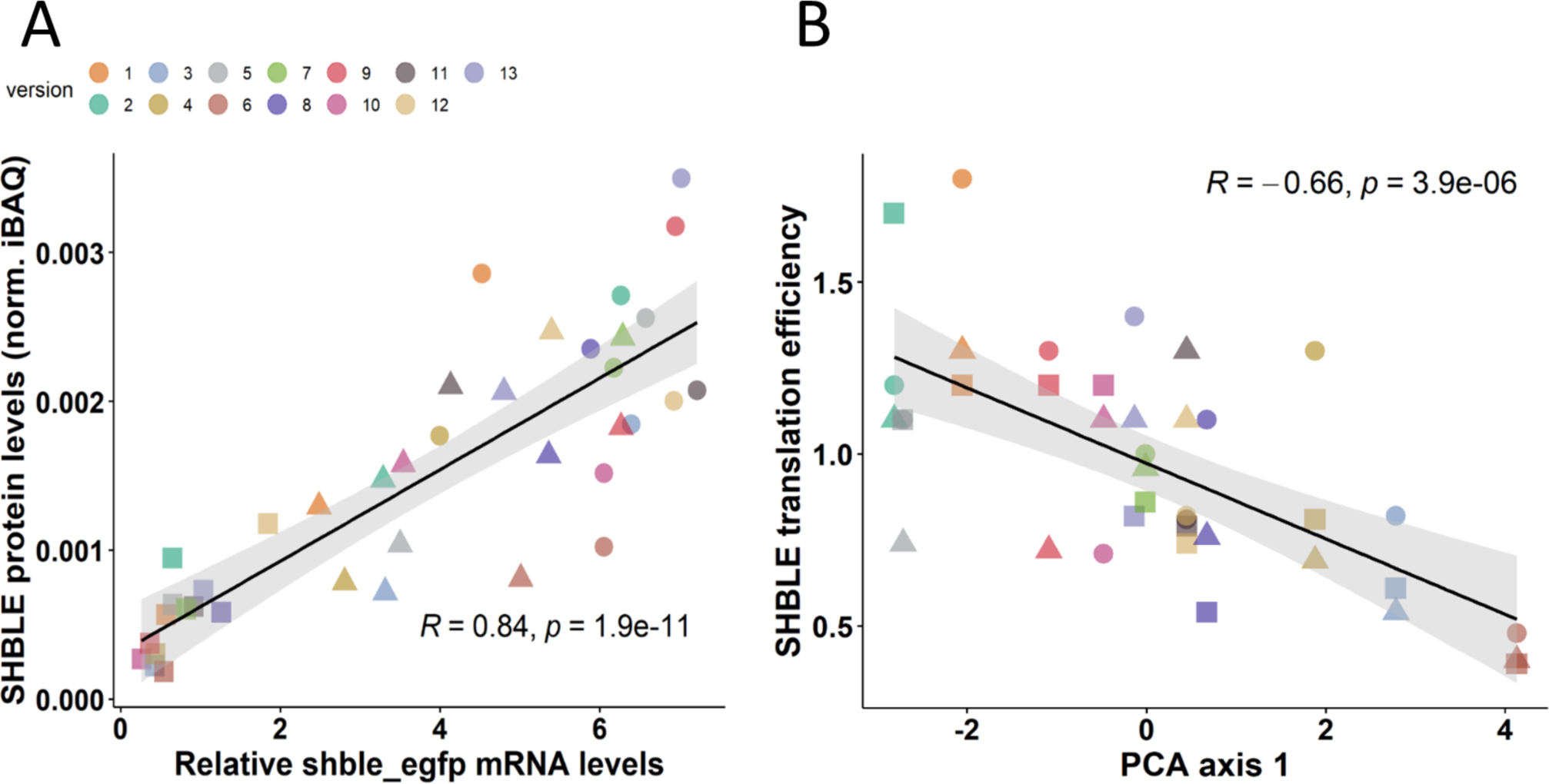
SHBLE production from synonymous versions of a bicistronic *shble_egfp* mRNA. (panel A) Pearson’s linear regression (black line) and 95% confidence interval of the fit (grey) between relative *shble_egfp* mRNA levels and relative SHBLE protein levels for the thirteen *shble* synonymous versions. **(panel B)** Pearson’s linear regression (black line) and 95% confidence interval of the fit (grey) between the projection on the first axis of PCA in Fig 1 for each *shble* version, and SHBLE translation efficiency. For both panel, values from a same biological replicate are represented by triangle, rectangle or circle shapes.

**Table 1:** Individual and sequential linear fits for variation of *shble* composition variables to account for variation of SHBLE translation efficiency. GC3, percentage of G or C at the third codon nucleotide codons; freq_CpG, CpG dinucleotide frequency; freq_TpA. TpA dinucleotide frequency; folding, energy of the more stable structure predicted for the mRNA shble ORF estimated with UNAfold online tool (http://unafold.org) (Markham et Zuker 2008); COUSIN_59, value of the COdon Usage Similarity Index of the shble version with respect to the average human codon usage (Bourret, Alizon, et Bravo 2019). AICc, corrected Akaike Information Criterion.

| Individual Fits |  |  |
| --- | --- | --- |
| Parameter | R square | AICc |
| GC3 | 0.464 | 5.13 |
| freq_TpA | 0.397 | 9.67 |
| freq_CpG | 0.22 | 19.73 |
| folding | 0.256 | 17.91 |
| COUSIN_59 | 0.396 | 9.74 |
| Sequential Fit |  |  |
| Parameter | R square | AICc |
| GC3 | 0.464 | 5.14 |
| COUSIN_59 | 0.482 | 6.32 |
| freq_CpG | 0.484 | 8.81 |
| freq_TpA | 0.486 | 11.44 |
| folding | 0.486 | 14.42 |
| Best fit | 0.464 | 5.14 |
| Sequential Fit (no GC3) |  |  |
| Parameter | R square | AICc |
| freq_TpA | 0.398 | 9.68 |
| COUSIN_59 | 0.435 | 9.67 |
| freq_CpG | 0.486 | 8.65 |
| folding | 0.486 | 11.45 |
| Best fit | 0.486 | 8.65 |

Since all the recoded *shble* versions are identical in the 5’ UTR and in the first 24 coding nucleotides, we understand that any differences in translation efficiency are related to translation elongation, rather than to translation initiation. Overall, our results demonstrate that synonymous recoding of the *shble* ORF is sufficient to impact SHBLE protein levels by affecting translation efficiency during translation elongation.

### Synonymous recoding of the upstream *shble* ORF does not affect downstream *egfp* ORF translation

Next, we applied the same analyses to explore whether variation in GFP protein levels could be accounted for by variation in mRNA levels and/or in mRNA composition. We used RT-qPCR and proteomic data as described above and included flow cytometry as an orthogonal technique to assess GFP levels. We integrated fluorescence intensity from a random sample of transfected cells (30,000 events) from each transfection experiment and used it as a proxy for GFP protein levels in the cell population. Our final dataset included fluorescence and mRNA levels evaluated on five replicates, additional to the three replicates in which GFP protein levels were assessed by proteomics. There was a very good correspondence between both GFP estimates, as variation in proteomic-based GFP levels accounted for 81% of the variation in cytometry-based GFP fluorescence levels (fig 5A, R^2^=0.83, *p=*2.1*10^-15^).

Variation in mRNA levels accounted for a significant fraction of variation in GFP levels, both using proteomic-based data (Fig5B, R^2^=0.55, *p=*7.7*10^-8^), and fluorescence-based data (Fig5C, R²=030, *p=*7.4*10^-8^). It is of note that the explanatory power of variation in mRNA levels on variation in protein levels is substantially higher for the upstream *shble* ORF, described above (Fig4A), than for the downstream *egfp* ORF. Translation efficiency of *egfp,* defined as GFP protein levels over *shble*_e*gfp* mRNA levels, was not different between *shble* synonymous conditions (Fig S5A and S5B, pairwise Wilcoxon rank sum test after B-H correction for multiple comparisons; α=0.05). This lack of differences on *egfp* translation efficiency among constructs remained true for both proteomic-based data, which displayed large experimental variation (Fig S5A, median value 0.84, range between 0.39 for *shble*#9 and 1.27 for *shble*#8), as well as for cytometry-based data, more homogeneous (Fig S5B, median value 0.81, range between 0.64 for *shble*#2 and 1.13 for *shble*#10). More importantly, *egfp* translation efficiency displayed no correlation with the overall compositional features of the bicistronic *shble_egfp* mRNA, when using both proteomic-based (Fig5D, R²=0.00, *p=*0.60) or cytometry-based data (Fig 5E; R²=0.02, *p=*0.17).

**Figure 5:**
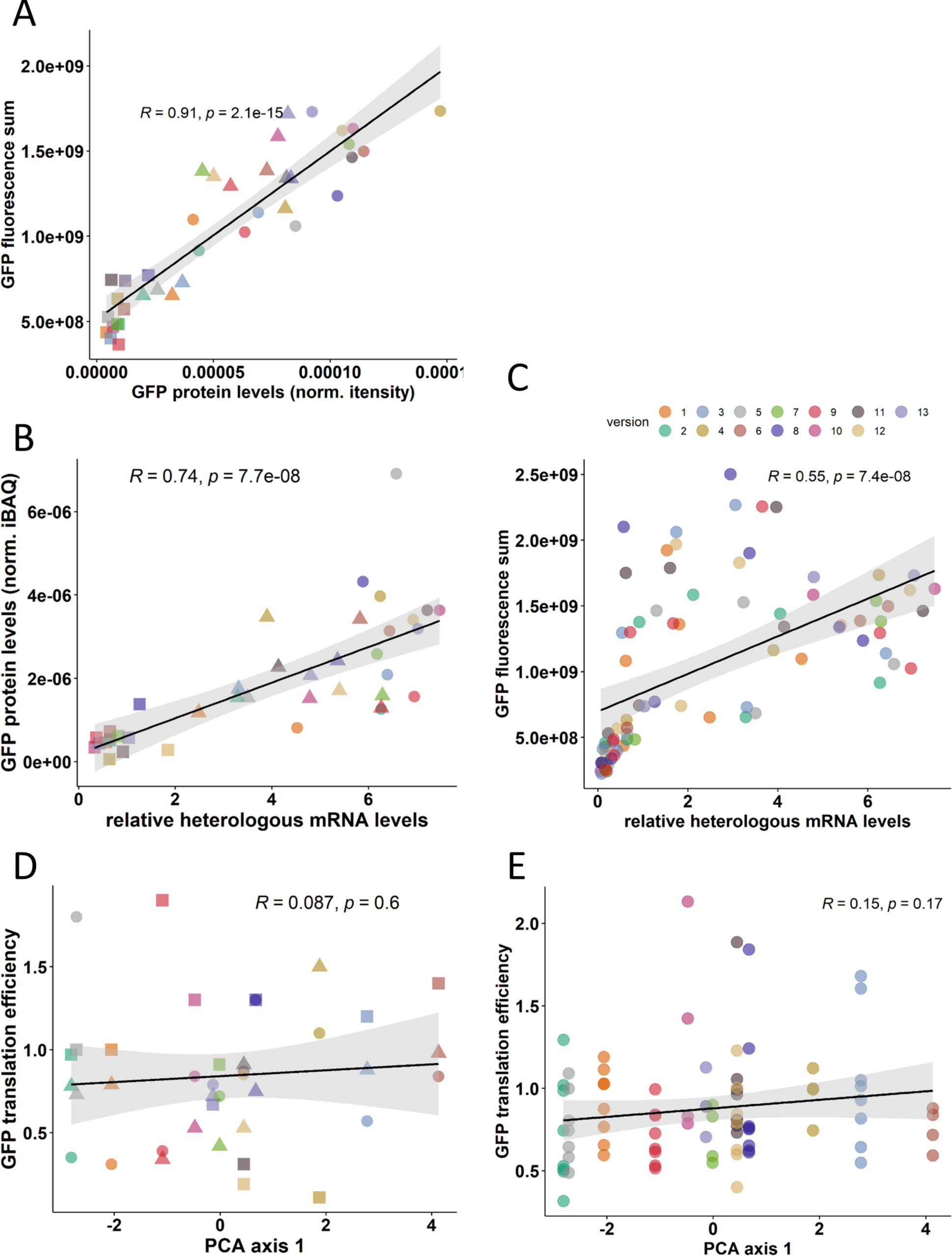
GFP production from synonymous versions of a bicistronic *shble_egfp* mRNA. (panel A) Pearson’s linear regression (black line) and 95% confidence interval of the fit (grey) between proteomic-based, sample-normalised GFP intensity levels and the sum of fluorescence intensity of the cellular population for the thirteen *shble* versions. Sums of fluorescence were calculated by integrating the fluorescence signal of 30,000 randomly selected cells in each treansfection event. **(panel B)** Pearson’s linear regression (black line) and 95% confidence interval of the fit (grey) between relative *shble_egfp* mRNA levels and relative GFP protein levels for the thirteen *shble* synonymous versions. Values from a same biological replicate are represented by triangle, rectangle or circle shapes. **(panel C)** Pearson’s linear regression (black line) and 95% confidence interval of the fit (grey) between relative *shble_egfp* mRNA levels and GFP fluorescence sum for the thirteen *shble* synonymous versions. **(panel D)** Pearson’s linear regression (black line) and 95% confidence interval of the fit (grey) between the projection on the first axis of PCA in Fig 1 for each *shble* version, and *egfp* translation efficiency calculated with proteomic data. Values from a same biological replicate are represented by triangle, rectangle or circle shapes. **(panel E)** Pearson’s linear regression (black line) and 95% confidence interval of the fit (grey) between the projection on the first axis of PCA in Fig 1 for each *shble* version, and *egfp* translation efficiency calculated with fluorescence data.

Overall, our results show that synonymous recoding of the upstream *shble* ORF does not have an impact on translation efficiency of the downstream *egfp* ORF.

### The downstream *egfp* ORF is probably translated by leaky scanning, independently of the nucleotide composition between ribosome binding site and *egfp* AUG

We have determined first the pervasive translation of the *egfp* downstream ORF, second the impact of the synonymous variation in the *shble* upstream on its own translation efficiency, and third the absence of impact of this synonymous variation in the upstream ORF the translation efficiency of the *egfp* downstream ORF. We sought then to identify the (non-)canonical mechanism(s) at play allowing the pervasive translation of a downstream ORF in eukaryotic cells. To do so, we have integrated results from mRNA levels and protein quantification for the upstream and the downstream ORFs.

We first evaluated the correlation between SHBLE and GFP protein levels, aiming at understanding whether ribosomal engagement in translating the upstream *shble* ORF was linearly accompanied by an engagement in translating the downstream *egfp* ORF. Globally, variation in SHBLE levels was only a moderate predictor of variation in GFP levels (Fig S6, R²=0.24; *p=*0.0017). Yet, when we stratified our data according to the match between the CUPrefs of the *shble* synonymously recoded ORF and those of the human average, a striking pattern appeared (Fig 6A): for *shble* versions overmatching the human average CUPrefs (*i.e. shble*#1, #2, #9, #12 and #13), SHBLE levels were in average 1,628 times higher than GFP levels (95%CI:1403-1854; R²=0.38, *p=*0.014), while for *shble* versions with CUPrefs close to the human average (*i.e. shbl*e#5, #7, #10 and #11) SHBLE levels were in average 998 times higher than GFP levels (95%CI: 831-1165; R²=0.55, *p=*0.0056), and for *shble* versions with CUPrefs under matching the human average (*i.e. shble*#3, #4, #6 and #8) SHBLE levels were in average 422 times higher than GFP (95%CI:394-503; R²=0.55, *p=*0.0058) (Fig 6A). The comparison of the three categories revealed significant increase in the SHBLE-over-GFP protein ratio following the increase in *shble* match to the average human CUPrefs (Fig 6B, pairwise Wilcoxon rank sum exact test with B-H correction; α=0.05). Finally, analysing SHBLE-over-GFP protein levels as a function of mRNA features further indicates that variation in the *shble* upstream ORF composition explains 30% of variation in SHBLE-over-GFP levels (R²=-0.30; *p=*0.00032) (Fig 6C). Finally, variation in GFP translation efficiency was not explained by variation in SHBLE translation efficiency, further suggesting a lack of influence of upstream ORF translation onto downstream ORF translation (Fig 6D, R²=0, *p=*0.64).

**Figure 6:**
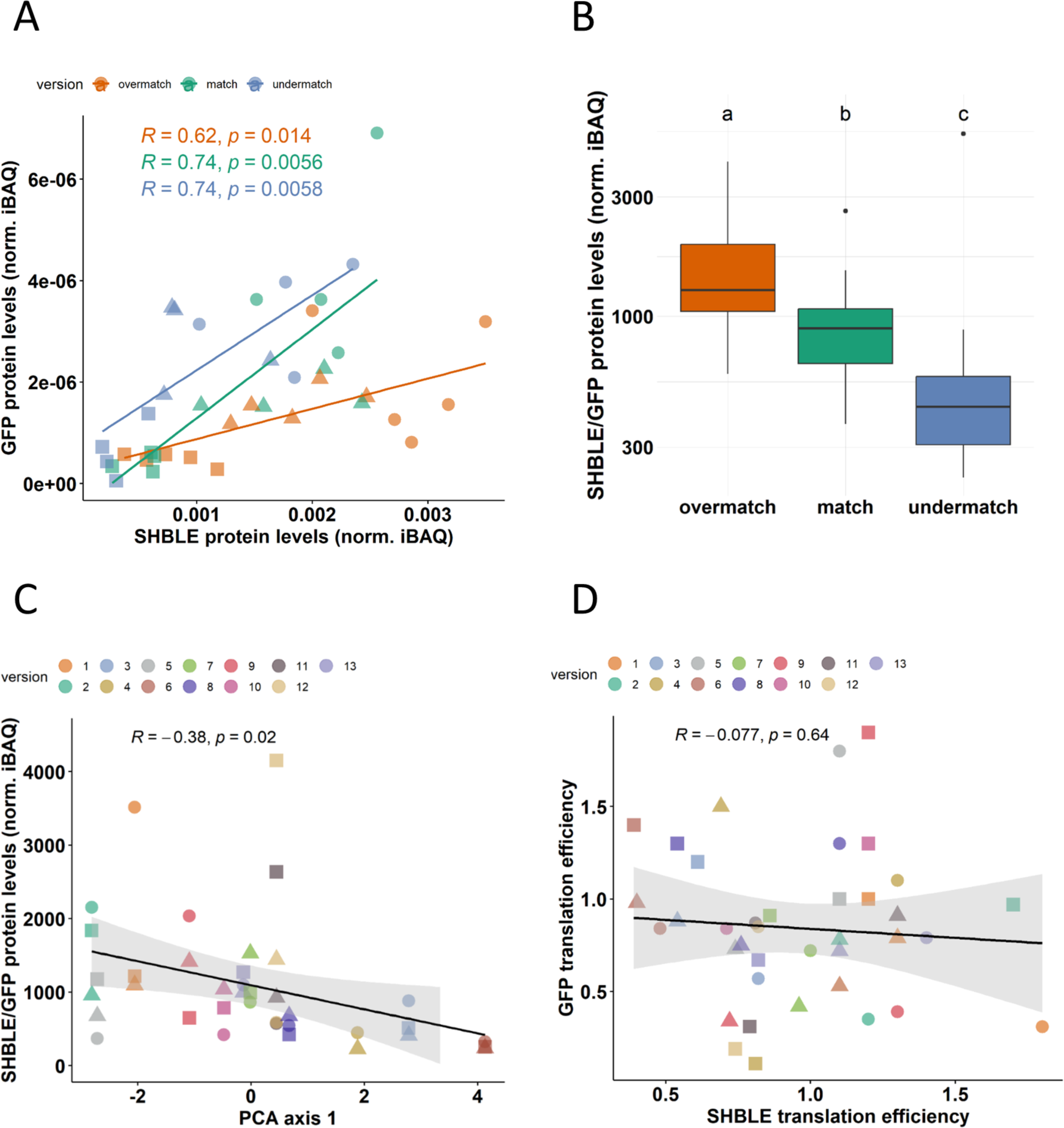
relative SHBLE and GFP production from bicistronic *shble_egfp* mRNA. (panel A) Pearson’s linear regression (coloured lines) between normalised SHBLE iBAQ levels and normalised GFP iBAQ levels for the thirteen *shble* versions, stratified by their match to the average CUPrefs of the human genome. **(panel B)** Box-and-whiskers plot of the SHBLE/GFP proteins ratio for the thirteen *shble* versions, stratified by their match to the average CUPrefs of the human genome. Letters present the results of a pairwise Wilcoxon rank sum test with B-H adjusted p-values; α=0.05; median values for samples labelled with the same letter are not statistically different. **(panel C)** Pearson’s linear regression (black line) and 95% confidence interval of the fit (grey) between the projection on the first axis of PCA in Fig 1 for each *shble* version, and the SHBLE/GFP proteins level ratios. **(panel D)** Pearson’s linear regression (black line) and 95% confidence interval of the fit (grey) between SHBLE translation efficiency and GFP translation efficiency. For panels A, C and D, values from a same biological replicate are represented by triangle, rectangle or circle shapes.

Overall, our protein quantification results show that synonymous variation in the *shble* upstream ORF modulates its own translation, but not that of the *egfp* downstream ORF. Given our experimental design with invariant 5’UTR, ribosome binding site and AUG translation context, we conclude that GFP is translated through infrequent events of ribosome leaky scanning which are not influenced by the nucleotide composition of the scanned sequence upstream of *egfp*’s AUG.

### Splicing events within the *shble* upstream ORF modulate both SHBLE and GFP protein levels but fall short to close the expression gap between the two

Five out of our thirteen *shble_egfp* engineered constructs produced bicistronic transcripts that undergo splicing events with donor and acceptor sites within the *shble* ORF. In these cases, the SHBLE protein can be produced only from the unspliced bicistronic mRNA, while the GFP protein can be conceptually produced from both the unspliced and spliced mRNAs, albeit with potentially modified translation efficiency. Two of these splicing events (*shble#4* and #6) had been described previously in (Picard et al. 2023) and correspond to the excision of an intron spanning nt 70 to nt 380 of the *shble* ORF (Fig 7A). This splicing event produces a *shble*I* ORF still in frame with the original *shble* and maintaining the same stop codon, thus conserving the 19-nt gap between the *shble* UAA stop codon and the *egfp* AUG start codon. The second splicing event concerns constructs *shble*#7, #10 and #13 and leads to the excision of an intron spanning nt 186 to nt 380 for *shble#7* and #10, or to nt 377 for *shble#13*. These three splicing events using the splice donor at nt 186 introduce a frameshift so that the <u>UGA</u> stop codon of the spliced, frameshifted *shble*II* ORF overlaps with the <u>AUG</u> start codon of *egfp,* within the CGC<u>AUGA</u>GC sequence.

**Figure 7:**
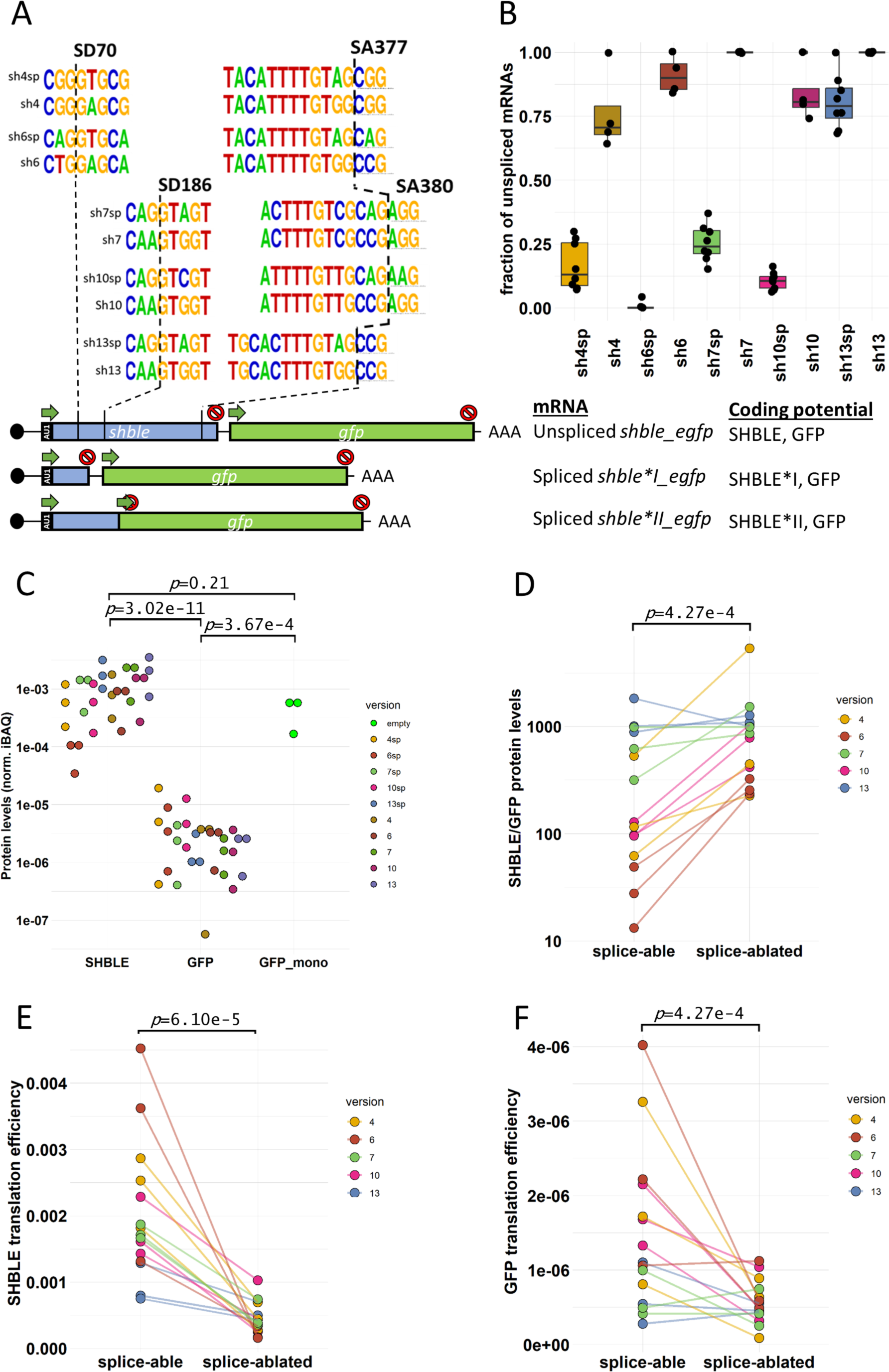
Effect of *shble_egfp* bicistronic mRNA splicing on SHBLE and GFP protein levels. (panel A) Schematic representation of unspliced and spliced bicistronic mRNAs generated for *shble* versions *shble*#4, #6, #7, #10 and #13 and their respective original or mutated splice site sequences. Figure should be read as follows, using *shble*#4 as an example: “sh4sp” refers to the sequence that undergoes splicing, while “sh4” refers to the sequence that has been mutated to ablate splicing. Splice donor (SD) and splice acceptor (SA) site position relative to the *shble* AUG are indicated by discontinuous lines. **(panel B)** Box-and-whiskers plot showing the fraction of unspliced mRNA generated by splice-able (*e.g.* “sh4sp”) and splice-ablated (*e.g.* “sh4”) constructs, determined by Bioanalyzer (see Fig S7 for details). **(panel C)** Dot plot showing relative levels of heterologous proteins measured by label free proteomic from three biological replicates. For each sample, iBAQ values of SHBLE and GFP were normalised by the total iBAQ value of the sample. The positive control “GFP_mono” condition translated GFP from a monocistronic *egfp* mRNA while the ten *shble* conditions translated SHBLE and GFP from a spliced or non-spliced bicistronic *shble_egfp* mRNA, as specified by the colour code. GFP values do not follow a normal distribution after a Shapiro normality test (*p* = 6.18e-13), hence the *p*-values present the probability that the medians of the groups are not different, after a pairwise Wilcoxon signed rank exact test (α=0.05). **(panel D)** Connected dot-plot showing SHBLE/GFP protein ratios for the ten splice-able and splice-ablated *shble* versions from three biological replicates. Paired differences between splice-able and splice-ablated versions were assessed using the Wilcoxon signed rank sum test (*p* = 4.27e-4). **(panel E)** Connected dot plot showing SHBLE translation efficiency from splice-able and splice-ablated pairs of constructs from three biological replicates. Paired differences between splice-able and splice-ablated versions were assessed using the Wilcoxon signed rank sum test (*p* = 6.10e-5). **(panel F)** Connected dot plot showing GFP translation efficiency from splice-able and splice-ablated pairs of constructs from three biological replicates. Paired differences between splice-able and splice-ablated versions were assessed using the Wilcoxon signed rank sum test (*p* = 4.27e-4).

In order to study the impact of splicing in the *shble* upstream ORF on the translation of the *egfp* downstream ORF, we introduced synonymous point mutations in each spliced *shble* version aiming at ablating the splicing capacity. All original and mutated donor and acceptor splice site sequences are displayed in figure 7A. Splicing efficiency was evaluated by means of bioanalyzer quantification (Fig S7A and B). Splicing efficiency of the original events ranged between 99% for *shble*#6 and 20% for *shble*#13 (Fig 7B). Mutational ablation drastically reduced splicing in all cases, allowing us to compare SHBLE and GFP translation (Fig. 7B). Assessment of total mRNA levels by RT-qPCR revealed no significant differences between constructs and no correlation between mRNA levels and splicing efficiency (Fig. S8A and B).

Both SHBLE and GFP proteins were detected in all samples from the ten constructs in biological triplicates, even those undergoing highly efficient splicing (Fig7C). SHBLE protein levels were on average 754 (95%CI: 581-927) times higher than GFP levels (Fig. 6C, Wilcoxon rank sum test with continuity correction; *p<*3.02*10^-11^). GFP protein levels produced from the *egfp* monocistronic control were not significantly different from SHBLE protein ones (Fig. 7C Wilcoxon rank sum exact test; *p=*0.2108) but most importantly, were significantly higher than GFP protein levels produced from the second ORF of every bicistronic mRNAs (Fig. 7C, Wilcoxon rank sum exact test; *p=*3.67*10^-4^). SHBLE protein was detected in all three label-free proteomic replicates for the *shble*#6sp condition (data in PRIDE entry PXD047576), albeit at the lowest levels across conditions (Fig 7C), even if we detected unspliced mRNA coding SHBLE in only one replicate out of eight (Fig 7B). This seems to indicate a better sensitivity of the proteomic measurements than of the transcriptomic ones following RT-PCR amplification and Bioanalyzer analyses. Mass spectrometry analyses did not detect any peptide that could correspond to the truncated SHBLE*I or SHBLE*II proteins, potentially translated from the corresponding spliced mRNAs. All our western-blotting efforts to detect these proteins by targeting the AU1-tag rendered also negative results (data not shown). Even if results in Fig 7C look qualitatively similar to results in figure 3, the slightly lower SHBLE levels and slightly higher GFP levels in the splice-able versions result in a significant decrease of the SHBLE-over-GFP protein levels due to splicing (Fig 7D, paired Wilcoxon signed rank sum test; *p* = 4.2*10^-4^). Specifically, splicing ablation resulted in SHBLE-over-GFP levels increase on average by 6.41 (CI95: 4.14-8.68), 11.61 (CI95: 8.14-15.08) and 6.87 (CI95:5.64-8.10) fold for *shble*#4, *shble*#6 and *shble*#10, respectively. In the case of *shble*#13, displaying low starting splicing levels, mutation of splicing signals did not result in a change of the SHBLE-over-GFP ratio (1.02 fold, CI95: 0.78-1.27). Finally, the response of *shble*#7 (2.40 fold, CI95: 1.25-3.55) was milder than those of *shble*#4, #6 and #10, although mutation totally ablated the high basal splicing activity of *shble#*7sp (Fig 7B, median efficiency 76%). We confirmed moreover that variation in splicing efficiency is a good predictor of variation in SHBLE-over-GFP protein ratio (Fig S9A, R²=-0.39, *p=*2.2*10^-4^). We obtained similar trends when estimating the impact of variation in splicing efficiency on variation in protein levels separately for SHBLE (Fig S9B, R²=-0.42, *p=*1.0*10-4), for proteomic-based GFP levels (Fig S9C, R²=0.29, *p=*2.1*10-3) and for fluorescence-based GFP levels (Fig S9D R²=0.42, *p=*1.4*10-8).

We next compared splice-able and splice-ablated versions in terms of translation efficiency. First, splicing ablation resulted in an increase in SHBLE protein levels (Fig 7C), as expected because splicing ablates the *shble* ORF. This increase in SHBLE levels was intriguingly accompanied by a decrease in SHBLE translation efficiency for a given *shble* version (Fig 7E, paired Wilcoxon signed rank sum test; *p*=6.1*10^-5^). The fact that we measured a very good linear covariation between *shble_egfp* mRNA levels and SHBLE protein levels (Fig S9F) does not support a possible saturation of the translation machinery for splice-ablated conditions. Comparison of splicing impact on SHBLE translation efficiency (Fig 7E) and on GFP translation efficiency (Fig 7F) reveals that: i) the drop in translation efficiency between splice-able and splice-ablated conditions is greater for SHBLE than for GFP; and that ii) for *shble#7* GFP translation efficiency is not impacted by splicing (Fig 7F and Fig S9E for cytometry data). Splicing resulted thus in an increase in GFP synthesis, accompanied by a significant increase in GFP translation efficiency (Fig 7F for proteomic data, paired Wilcoxon signed rank test; *p*=4.27*10^-4^, and Fig S9E for cytometry data, paired Wilcoxon signed rank test, χξ=0.05).

Overall, our results show that splicing in the upstream *shble* ORF results in a qualitative increase in translation of the downstream *egfp* ORF. Nevertheless, the quantitative extent of this increase depends on the specific synonymous recoding of each *shble* version.

## Discussion

In this work, we present a straightforward experimental model, composed by synonymous versions of a short *shble* upstream ORF followed by an *egfp* downstream reporter ORF, conceived to study gene expression regulation from virus-like bicistronic mRNAs in human cells and to allow for mRNA and protein quantification. Our results show first that the human cellular translation machinery can indeed translate the downstream ORF in bicistronic mRNAs, independently of the characteristics of the upstream ORF. Even if protein levels translated from the downstream ORF are hundreds of times lower than those from the upstream ORF, our results openly challenge the mainstream textbook interpretation positing that the eukaryotic translation machinery does not handle the downstream ORFs in bicistronic transcripts. Second, we show that translation efficiency during elongation of the upstream ORF is largely determined by its own nucleotide composition. Third, the upstream nucleotide composition does not influence translation of the downstream ORF, but splicing within the upstream ORF leads to an increase of downstream ORF translation.

Two main mechanistic scenarios can be invoked to account for GFP synthesis in our experimental system: i) leaky scanning, *i.e.* the recruited ribosome does not engage in translation of the upstream ORF and translates instead the downstream ORF; and ii) translation reinitiation, *i.e.* the ribosome remains engaged in translation and the downstream ORF is translated immediately upon termination of upstream ORF translation. By design, we anticipate that ribosomal recruitment and engagement do not differ among constructs, as the 5’UTR and the first 24 coding nucleotides of the upstream ORF are identical. Differences in translation efficiency of the upstream ORF among constructs will thus arise essentially during the elongation phase. Also by design, the context between the two ORFs is shared across constructs as the last 24 coding nucleotides of the upstream ORF, its stop codon, the 19 nucleotides spacer and the full downstream ORF are identical for all versions. Differences among constructs in GFP synthesis will thus be linked to a differential probability for leaky scanning and/or by a differential probability for translation reinitiation upon termination, arising as a function of the compositional features and splicing of the upstream *shble* ORF. We have applied this reasoning for discussing our results.

### Synonymous variations in the *shble* upstream ORF introduce cryptic splicing events but does not affect heterologous mRNA levels

Our previous study identified splicing events in *shble*#4 and *shble*#6 (Picard et al. 2023). From the nine newly tested synonymous *shble* versions, three harboured functional splice sites, namely *shble*#7, *shble*#10 and *shble*#13, recognized with varying efficiency degrees by the U-2 OS human cell model: splicing was highly efficient for *shble*#4, #6, #7 and #10, with less than 25% unspliced mRNAs while it was inefficient for *shble*#13, with above 80% unspliced mRNA. Splicing of *shble*#4 and #6 mRNAs was even more efficient in U-2 OS cells than in HEK293 cells (84% *vs* 21% for *shble*#4 and 99% *vs* 79% for *shble*#6 in U-2 OS compared to data reported for HEK293 (Picard et al. 2023)). All splice events occurred within the *shble* ORF, thus ablating the SHBLE coding potential in the spliced mRNAs. The donor splice sites resembled the second most common among the typical U1 and U2 splice sites (45.1% of the splice sites in human transcripts) (Sibley, Blazquez, et Ule 2016). In contrast, while the intron 3’ sequences resembled also the typical U1 and U2 sites, the 5’ of the downstream exons did not match any of the common typical or atypical U1 and U2 sites nor the typical U11 and U12 sites (Sibley, Blazquez, et Ule 2016). This is probably the reason for which none of these splice events were confidently predicted by commonly used algorithms (Solovyev 2003; Desmet et al. 2009). We introduced synonymous mutations in all five acceptor and donor sites, modifying two to four nucleotides, aiming at ablating splicing (Fig.7A). We tried to diverge from the typical splice signal, *i.e.* removing the GT and AG dinucleotides at the 5’ and 3’ of the intron, respectively (Eskesen, Eskesen, et Ruvinsky 2004). We were nevertheless constrained by the need to conserve the synonymous coding *shble* sequence and reading frame, so that for instance mutated *shble*#7, *shble*#10 and *shble*#13 still retained the consensus GT dinucleotide at the intron 5’ end. Overall, splice site mutations resulted in total ablation of splicing in *shble*#7 and *shble*#13, and in substantial reduction of splicing in all other versions, allowing us to compare splice-able and splice-ablated pairs of constructs. A residual splicing activity around 20% remained for *shble*#4. Most likely, remaining branchpoint sequences, polypyrimidine tract in the intron 3’ end, as well as splice enhancer *cis*-elements may still suffice for providing the observed splicing activity (K. Gao et al. 2008; Sibley, Blazquez, et Ule 2016; Wang et Burge 2008; Fallot et al. 2009). Our results suggest that cryptic splicing is conspicuous after synonymous recoding, as it seems to be the case for many natural transcripts in eukaryotes (Bénitière, Necsulea, et Duret 2024), as we detected it in five out of the thirteen synonymous variants, and splicing efficiency may eventually depend on the presence/absence of a limited number of local nucleotide changes.

CUPrefs have an impact on mRNA production and stability (Hanson et Coller 2018). We have here experimentally addressed the impact of codon recoding on heterologous mRNA levels. In our system, neither synonymous sequence recoding nor splicing led to differences in terms of total heterologous mRNA levels. Our results are in first view in contradiction with data from other eukaryotic experimental systems, such as *Neurospora* (F. Zhao et al. 2021; Zhou et al. 2016), *Saccharomyces* (Victor et al. 2019; S. Chen et al. 2017; Presnyak et al. 2015), or human cells (Newman et al. 2016), which have shown that CUPrefs are an important determinant of steady-state mRNA levels. It has been proposed that this effect is mediated by variation in nucleotide composition and CUPrefs in the transcript 5’ end (S. Chen et al. 2017). It has also been proposed that the impact of CUPrefs on transcription is promoter-dependent (Yang et al. 2021). The lack of impact on mRNA levels in our system could thus be explained by our choice of experimental model. All our constructs share identical 5’UTR and first 24 coding nucleotides in the *shble* ORF, as well as identical 3’UTR *cis*-regulatory sequences and poly-A signals, which have also been proposed to regulate transcript half-life (Cheng et al. 2017). The overall invariant mRNA steady levels in our system are thus consistent with a low impact of variation in nucleotide composition and mRNA folding energy in the coding portion of the bicistronic mRNAs and most importantly no variation of the 5’ and 3’ UTR of mRNAs.

### Compositional features of the *shble* upstream ORF modulate its own translation efficiency during elongation

We have engineered 13 synonymous *shble* versions to study the impact of upstream ORF composition on its own translation and on the translation of the downstream *egfp* ORF. In our design, the molecular contexts for *shble* translation initiation and translation termination are strictly identical between versions. Our results show that variation in composition variables of *shble* elongation phase is sufficient to modulate its own translation, so that *shble* versions overmatched to the average human CUPrefs are more efficiently translated than human under-matched *shble* versions.

To explore the impact of synonymous recoding on bicistronic mRNAs translatability we have addressed protein-over-mRNA levels, as a proxy to study translation efficiency (Hernandez-Alias et al. 2023). Our results show that 44% of the variation in *shble* translation efficiency is explained by variation in *shble* nucleotide composition: the stronger the bias towards human-average CUPrefs of the *shble* version, the higher the *shble* translation efficiency (fig 4B and S4). Maximum differences in SHBLE translation efficiency across constructs were around 3.2-fold between human over-matched *shble*#1 and human under-matched *shble*#6. Such magnitude in translation efficiency differences associated to nucleotide composition and CUPrefs can appear modest compared to literature reporting differences up to 100-fold in protein levels between synonymous variants of a same ORF in different eukaryotic systems, *e.g.* yeast (Kaishima et al. 2016), mammalian cells (Nagata et al. 1999) or plant cells (Kwon et al. 2016). However, the fact that in our experimental system the translation initiation context remains invariable across all *shble* versions is consistent with such moderate effect, in agreement with the most cogent and most comprehensive synonymous recoding study to date. Indeed, even after having synthesised and explored over 240,000 synonymous variants in a large endeavour to understand the impact of nucleotide composition on prokaryotic translation, the beautiful and exhaustive experimental plan and statistical analysis by Cambray and co-workers could only explain 30% of the total variance in protein production. These authors identified initiation of protein synthesis as the most determinant step for total protein production. Variation in the folding energy of the mRNA structure(s) spanning 30 nucleotides upstream and downstream the AUG start codon, which corresponds to the average ribosomal mRNA occupancy (Ingolia, Lareau, et Weissman 2011), accounted alone for around 25% of the total variation in protein production (Cambray, Guimaraes, et Arkin 2018). Our experimental design however, strictly focuses on differences in translation elongation, and we consider thus that the mRNA structure in the vicinity of the *shble* AUG start codon is essentially the same across constructs, and does not come in to play. As such, we interpret that the impact we observe from variation in CUPrefs on variation in SHBLE production efficiency is linked directly to translation elongation. Our results using human cells are consistent with previous experimental data showing that synonymous codon recoding can affect eukaryotic translation elongation respectively in *Neurospora, Drosophila* and yeast (Yu et al. 2015; F. Zhao, Yu, et Liu 2017; Pop et al. 2014). The reported modulation of translation efficiency at constant translation initiation supports further the claim that translation elongation can influence translation initiation (Chu et al. 2014; Lyu et al. 2021).

The impact of the so-called “gene optimization” on protein expression is such that a growing number of algorithms have been published (Sandhu et al. 2008; Chin, Chung, et Lee 2014; Fu et al. 2020) describing different engineering approaches for tuning composition variables to maximize mRNA, protein and protein-to-mRNA production for heterologous expression in biotechnology (F. Gao et al. 2003; Graf, Deml, et Wagner 2003; Nogales et al. 2014; Broadbent et al. 2016); for a review see (Mauro and Chappell 2014). However, the literature shows that the ability to manipulate and predict the behaviour of the translation machinery does not necessarily imply understanding it. In our experimental system, after having established that variation in the overall biochemical features of the *shble* synonymous variants explains 44% of variation in its own translation efficiency, we aimed at disentangling the individual impact of compositional variables.

In our bicistronic system, variation in GC3 composition in the *shble* upstream ORF explained 46% of the variation in SHBLE production efficiency, in good accord with our previous study using *shble*-*P2A*-*egfp* monocistronic expression system (Picard et al. 2023). The large explanatory power of *shble* GC3 composition on translation efficiency fits also well the results communicated for intronless reporter genes (Mordstein et al. 2020). The individual explanatory power from CUPrefs, TpA or CpG frequency, and mRNA folding energy was already accounted for by the explanatory power provided by variation in GC3. Only when variation in GC3 was not considered, combined variation in TpA frequency, in CpG frequency and in CUPrefs accounted for 48% of the variation in *shble* translation efficiency (Table 1). Even in this case, the introduction of variation in mRNA folding energy did not increase the explanatory power over SHBLE production efficiency. Our results support the view that mRNA folding downstream the AUG start codon context influences only marginally translation efficiency, in agreement with the well-established literature showing that effects of mRNA folding energy on protein levels are exerted mostly on translation initiation (Kudla et al. 2009; Boël et al. 2016; Cambray, Guimaraes, et Arkin 2018).

Dinucleotide frequencies and their impact on different steps of gene expression are particularly interesting, and underrepresentation of CpG and TpA dinucleotides across a broad panel of genomes is a well-established observation (Swartz, Trautner, et Kornberg 1962; Beutler et al. 1989; Simmonds et al. 2013). CpG dinucleotides are under negative selective pressure in eukaryotes as they negatively regulate mRNA life: at the DNA level, intragenic CpG methylation is a potent repressor of transcription (Bauer et al. 2010; Kosovac et al. 2011; Radrizzani et al. 2024); at the RNA level, CpG-rich transcripts can be targeted for degradation, through a mechanism proposed to recognize non-self mRNAs by endonucleases (Takata et al. 2017; Duan et Antezana 2003). Regarding TpA dinucleotides, their underrepresentation in coding sequences is hypothesised to be selected for as it alleviates the risk of deleterious nonsense mutations (Beutler et al. 1989), as well as it prevents mRNA recognition by endonucleases (Odon et al. 2019). CpG frequency is highly correlated to total GC content and their respective effects on translation are consequently hard to disentangle (Mordstein et al. 2020). Still, the high CpG frequency of *shble*#5 in our system seemed to have a negative impact on its own translation efficiency compared to constructs with similar GC3 content, like *shble*#1 and *shble*#2 (Fig S4B). As *shble* sequence recoding showed no impact on mRNA levels across constructs, we interpret that extremely high CpG content could negatively impact protein levels, directly by modulating translation. On the contrary, TpA frequency is inversely correlated to GC content and thus to *shble* translation efficiency by human cells (Fig S4E). As no construct harboured varying TpA frequency with similar GC content, we cannot conclude on a possible effect on translation for the range of TpA frequency tested.

Finally, in our study we have not considered the impact of codon pairs on translation efficiency. Intuitively, the functional translation unit in the ribosome is rather the di-codon, as it can be conceived that the chemistry of the codon-tRNA complex in the P site may alter the codon-anticodon recognition chemistry for tRNAs entering the A site. The impact of dicodon frequency on translation is well documented in global transcriptome-proteome studies (Alexaki et al. 2019; Doyle et al. 2016), as well as in synonymous recoding, mostly from virus models (Conrad et al. 2018; Eldemery et al. 2023). However, this variable is often overlooked in experimental studies, even in those with much larger scale than ours (Cambray, Guimaraes, et Arkin 2018), essentially for two reasons: first, because the large di-codon combinatory space requires strong effects for pinpointing statistically significant differences; and second because it is intrinsically complicated to disentangle the effect of di-codons from that of di-nucleotides over the codon-codon boundary (Kunec et Osterrieder 2016; Daron et Bravo 2021; Alonso et Diambra 2023).

### The bicistronic mRNA organization is a major factor conditioning protein synthesis in human cells

Biochemistry textbook descriptions of translation emphasize that eukaryotic mRNAs are monocistronic, and that the eukaryotic machinery cannot handle the downstream ORF located in bicistronic mRNAs, as opposed to prokaryotes. In contrast to this standard view, our results show that human translation machinery can translate the downstream ORF located in *shble-egfp* bicistronic mRNAs, even if the resulting GFP protein levels are about over thousand times lower than SHBLE protein levels.

Experimental work had already demonstrated in the 1980s the translation of a downstream ORF from a bicistronic mRNA in human cells (Subramani, Mulligan, et Berg 1981; Mertz, Murphy, et Barkan 1983). Further pioneering works on the regulation of translation initiation established that adding an out-of-frame AUG codon upstream an ORF impaired its translation, but that adding a stop codon between the two ORFs and in frame with the first ORF, thence creating what is called now a short upstream ORF, rescues translation from the downstream ORF (Dixon and Hohn, 1984; Hughes et al., 1983; Kozak, 1984; Liu et al., 1984). Subsequent work using biochemical approaches to quantify downstream ORF translation from bicistronic mRNAs determined drops of between 5 to 10 fold and 100 to 300 fold, compared to translation of the same ORF from monocistronic mRNA (D S Peabody et Berg 1986; David S. Peabody, Subramani, et Berg 1986; Kaufman, Murtha, et Davies 1987). The currently consolidated view is that in eukaryotes, the presence of an upstream ORFs can modulate translation efficiency of downstream ORFs (Hellens et al. 2016; Hinnebusch, Ivanov, et Sonenberg 2016) and it is commonly understood that both the presence of introns in the 5’UTR and the presence of an upstream ORF result in a decreased translation of the downstream ORF (Lim et al. 2018). Nevertheless, in a systematic exploration of 4096 variants of a 21-nucleotides long upstream ORF with synonymous differences in three codons, Lin and coworkers reported effects from two-fold increase to two-fold decrease in translation of the downstream ORF (Lin et al. 2019).

Our study qualitatively confirms that human cells can translate upstream ORFs and downstream ORFs located onto the same bicistronic mRNAs, and provides further quantitative evidence of the enormous impact of ORF position in translation engagement. We have used normalised iBAQ data obtained from label-free proteomics to measure protein levels, and we estimate SHBLE levels to be in average over 1197 times higher than the accompanying GFP levels (fig 3). Notwithstanding, these results do not mean that in our cells there is one GFP molecule for every 1197 SHBLE molecules. While label-free proteomic results allow comparing levels between two different proteins, it is essential to keep in mind that different proteins display different responses across the full procedure of digestion, fragmentation, peptide separation, detection and identification, and thus different detectability (L. Zhao et al. 2020; Arike et al. 2012). Indeed, our previous experiment using a *shble*-*P2A*-*egfp* monocistronic expression system, in which SHBLE and GFP were translated in equi-molarity thanks to the ribosomal skipping step introduced by the P2A signal (Liu et al. 2017), label-free proteomics still revealed a 1:2 detection ratio for SHBLE:GFP proteins (Picard et al. 2023). This suggest that label-free proteomic data could have even under estimated by a factor of two the actual SHBLE-over-GFP levels.

Our results show that increased SHBLE levels are accompanied of increased GFP levels in all cases (Fig 6A). This covariation was not homogeneous among synonymous *shble* versions, as SHBLE-over-GFP ratios varied with the match between CUPrefs of the *shble* ORF and the human average ones (Fig 6B), and differed 7.1 times between the highest values for human over-matched *shble*#1 and the lowest values for human under-matched *shble*#6. Overall, 30% of variation in SHBLE-over-GFP ratio across constructs was accounted for by variation in the *shble* compositional features (Fig S6C). Thence, any mechanistic interpretation of EGFP production in our system will need to explain that: i) *shble* translation efficiency varies as a function of its own composition (Fig 4A); ii) that this is not the case for *egfp* (Fig 6B); and iii) that there is no covariation between the translation efficiency values for the two ORFs (Fig 6B). Appealing to the explanation of leaky scanning, we would interpret from our results that ribosomal engagement on translating the upstream ORF is more efficient in human-matched coding sequences, so that for a similar mRNA ability at recruiting ribosomes and for similar mRNA levels, higher levels of SHBLE synthesis would be accompanied by lower levels of GFP synthesis, resulting in the observed increased SHBLE-over-GFP ratio for human-matched *shble* versions. This interpretation would nevertheless imply that ribosomal engagement on the *shble* ATG is modulated by a downstream mRNA region, not covered by the ribosome. Appealing to the explanation of reinitiation upon termination, we would interpret that nucleotide composition of the *shble* coding region has a direct impact on downstream ORF translation, so that termination efficiency is higher and/or the probability of ribosomal re-engagement is lower in human-matched coding sequences. In its turn, this interpretation would imply that the termination process is modulated by an upstream mRNA region, not covered by the ribosome. We have resorted to the comparisons of the paired splice-able and splice-unable versions in our system to identify the differential experimental support for these two alternative explanations.

### Splicing in the *shble* upstream ORF modulates both SHBLE and GFP protein levels but fall short to close the expression gap between the two

In eukaryotic transcription, the presence of introns leads to increased mRNA levels, by a well-established but not fully understood mechanism called intron-mediated enhancement (Callis, Fromm, et Walbot 1987; Brinster et al. 1988). Indeed, AU-rich mRNAs can be retained in the nucleus or in specific cytoplasmic structures such as P-bodies and may not be available for translation, thus resulting in lower protein levels and decreased translation efficiency when normalising by total mRNA levels (Courel et al. 2019; Mordstein et al. 2020). Splicing can also enhance translation, as intronless transcripts result in lower protein levels than their counterparts containing an intron in the 5’UTR (Matsumoto, Wassarman, et Wolffe 1998; Mordstein et al. 2020). This effect is again mainly mediated by modulation of mRNA subcellular location (Le Hir, Nott, et Moore 2003). Indeed, spliced mRNAs are more efficiently exported from the nucleus than identical, intronless transcripts (Luo et Reed 1999). Subcellular location has been shown to largely determine translatability in eukaryotes (Le Hir, Nott, et Moore 2003). In our experimental setup, we did not observe differences in total heterologous mRNA levels neither among constructs in general, nor between pairs of mRNA constructs solely differing in their ability to undergo splicing (Fig S8A). In contrast, we observed a systematic increase in GFP translation efficiency in constructs that undergo splicing compared to those that do not, and this increase was significant for *shble*#4, #6 and #10 (Fig 7F). Given that GFP can be synthesised from both the spliced and the unspliced constructs, we interpret that this increase in GFP translation efficiency is related to an increased translatability of the *shble*_egfp* spliced mRNA. The presence of splicing signals in the upstream *shble* ORF could thus positively regulate translation of the downstream *egfp* ORF, by depositing regulatory protein complex on splice junction which in turn facilitate nuclear export and subsequent availability for the translation machinery (Le Hir, Nott, et Moore 2003).

The splice events detected in our system are not homogeneous with regards to the C-terminus of the SHBLE*I and SHBLE*II proteins. While the splicing for *shble*#4 and #6 conserves the *shble* stop codon, the 19bp intergenic and the *egfp* start, the splicing event in *shble*#7, #10 and #13 causes a frameshift that places the UGA *shble*II* stop codon overlapping the AUG *egfp* start codon. A similar <u>A*UG*</u>*A* configuration has evolved in Feline calicivirus to facilitate translation reinitiation, but it is strongly dependent on a specific mRNA secondary structure (Powell, Brown, et Brierley 2008). It could be thus claimed that a cryptic signal for initiation upon termination uncovered by the unpredicted splice events allows for GFP synthesis from the *shble*#7, #10 and #13 spliced mRNAs. However, the efficiency of this mechanism strongly depends on a very specific mRNA secondary structure (Pöyry et al. 2007), and it would still not explain GFP synthesis from *shble*#4 and #6, which would need a second, not yet described termination-reinitiation signal.

Very interestingly, this increase in GFP levels was not accompanied by the presence of the SHBLE*I and SHBLE*II proteins, corresponding to the modified *shble* ORFs upon splicing that could potentially be synthesised from the spliced versions *shble*#4 and #6 and from *shble*#7, #10 and #13. Indeed, we found no peptide that could have originated from any of these proteins in our label-free proteomic assays. We should first raise word of caution, because the N-terminus of these frameshifted proteins is identical to SHBLE, and the number of peptides that could allow for differential identification is very restricted. It could further be argued that these protein forms could be difficult to detect by mass spectrometry, as they are short and could have been produced at very low levels (Wacholder et Carvunis 2023). They remained nevertheless undetected after numerous western-blot attempts, under the same conditions (tag and antibodies) that allowed detection of the full-length SHBLE protein. We interpret from our data that the amount of the SHBLE*I and SHBLE*II proteins translated from the upstream ORF in the spliced mRNAs is negligeable. Should this be the case, the increase in GFP production from the spliced transcripts could obviously not be due to translation reinitiation upon termination, and we would thus claim leaky scanning as a better explanatory hypothesis for our observations. This interpretation is further supported by the descriptions showing that spliced bicistronic mRNAs are enriched in polysomes fractions compared to unspliced mRNAs (Fallot et al. 2009), and explaining the significantly increased levels of *egfp* translation efficiency for *shble*#4, #6 and #10 in the absence of changes in total mRNA levels.

## CONCLUSION

We have conceived and tested a toy model system to study translation from a bicistronic mRNA in human cells. Our results support the view that mRNA nucleotide composition and synonymous codon substitutions can have a direct impact on translation efficiency via differential codon-anticodon interactions, but that they further modulate protein synthesis by the introduction/removal of splicing signals and mid-and long-range intramolecular interactions (Callens et al. 2021). We show that modification of CUPrefs exclusively during elongation modifies overall translation efficiency and that it can result in the appearance of cryptic, unpredicted splice signals. We demonstrate that the ORF located downstream in a bicistronic mRNA can be translated by human cells, albeit at much lower levels than the upstream ORF. We finally propose that our experimental results about the impact of variation in CUPrefs and splicing in the upstream ORF on the translation of the downstream ORF are largely compatible with a mechanism of leaky scanning. We anticipate that our results will help understand the global role of upstream ORFs in the eukaryotic genomes. Finally, we specifically hope that our work will help shed light onto the mechanisms by which viruses manipulate the eukaryotic translation machinery, resulting in protein synthesis from complex, polycistronic viral transcripts.

### Experimental procedures

#### Design of the *shble* synonymous versions and plasmid constructs

Thirteen synonymous versions of the *shble* gene were designed to explore the impact of open reading frame composition variables on gene expression (all sequences available on GenBank, see data availability paragraph). As previously described in (Picard et al. 2023), versions *shble*#1 to #6 were designed applying the “one amino acid - one codon” approach, *i.e.*, all instances of a given amino acid were recoded with the same codon, depending on their frequency in the human genome and the GC-rich or AT-rich choice. For versions *shble*#7 to #13 we designed first, a thousand “guided random” synonymous *shble* sequences with CUPrefs based on the average human ones (Puigbo et al. 2007) and we chose seven based on their match to the human CUPrefs as evaluated by the codon adaptation index (Sharp et Li 1987) and on their mRNA folding energy values of the *shble* sequence, calculated using UNAfold online tool (http://unafold.org) (Markham et Zuker 2008). Identical nucleotide sequences encoding for a N-terminal AU1 tag (amino acid sequence MDTYRI), a C-terminal FLAG tag (amino acid sequence DYKDDDDK) and a TAA a STOP codon were added to all *shble* versions. All 13 versions were chemically synthesised (GenScript), cloned in the pCDNA3.1(+)-EGFP-C (Invitrogen) expression vector on the Xho1 restriction site. Schematic depiction of vector organization is displayed in Fig S1A. For all sequences, we calculated nucleotide composition variables GC percentage in the third codon position (GC3), CpG dinucleotide frequency, TpA dinucleotide frequency, CUPrefs to the human average using the COUSIN online tool (http://cousin.ird.fr) (Bourret, Alizon, et Bravo 2019) and mRNA folding energy of the *5’UTR*-*au1-shble* portion (TABLE S1).

#### Splicing discovery, experimental design and data analyses

Our previous study on *shble*#1 to #6 showed that transcripts from *shble*#4 and *shble*#6 underwent splicing (Picard et al. 2023). In the preliminary results for the present study, we identified by means of RT-PCR and Sanger sequencing three additional *shble* versions subject to splicing with varying efficiency, namely *shble*#7, #10 and #13. We introduced thus synonymous changes in both donor and acceptor splicing sites for all five constructs aiming at ablating splicing. Modified sequences are described in Fig 7A and S2C. Vector information and complete plasmid sequences are available on GenBank (see data availability paragraph).

#### Cell culture and transient transfection

The U-2 OS cell line (ATCC, HTB-96) was cultured at 37°C with 5% CO2 in McCoy’s 5A medium supplemented with L-glutamine (Biowest), with 10% heat-inactivated foetal calf serum (FCS, EuroBio) and with 1% penicillin-streptomycin (Fisher scientific). Transient transfection was performed using Turbofect according to the manufacturer’s instructions (Thermo Scientific). One million cells were seeded in a T25 flask 24h prior to transfection. Cells were transfected with a mix of 12µL Turbofect (Thermo Fisher Scientific) and 3 µg of the corresponding vector in 2% FBS McCoy’s 5A for 6h, and then incubated in McCoy’s 5A 10% FCS. Cells were harvested 24h after transfection start. For all replicates we used as transfection controls the pcDNA3.1(+)-C-EGFP (designated as “empty”), which expresses a monocistronic *egfp* mRNA, and a mock transfection using exposing the cells to the transfection agent alone, without heterologous DNA (designated as “mock”).

#### Cell collection for downstream experiments

Briefly, 24h after transfection cells were washed in Ca^2+^/Mg^2+^-depleted PBS (Biowest) and treated with trypsin 0.25%, 0.53mM EDTA in Ca^2+^/Mg^2+^-depleted PBS (Biowest) for 5 minutes. Cells were resuspended in cold McCoy’s 5A medium and split into four aliquots: one for cytometry, one for RNA extraction and two for protein extraction. Both RNA and protein cell pellets were stored at -70 °C until further use.

#### Flow Cytometry

Pelleted cells were fixed in cold PBS, 2% paraformaldehyde (Sigma) for 15min, washed twice in cold PBS 0.1M glycine (Sigma) and analysed within 24h. Flow cytometry was performed at the MRI imaging facility (Montpellier, France) on a NovoCyte flow cytometer system (ACEA biosciences) running the NovoExpress software (v1.6.2) with the following measurement settings: a hundred thousand ungated events at fast flow rate (approx. 2,000 to 3,000 events/s) with FSC.H superior to 100.0 arbitrary units (A.U.). GFP fluorescence was acquired by excitation at 488 nm with PMT gain set to 479. Filtering of cell debris and doublets, and downstream cell population analyses were done with an in-house R script available on git-hub (see data avaibility paragraph).

#### Qualitative and quantitative analyses of heterologous mRNA

Total RNA was extracted using the Monarch total RNA miniprep kit (NEB). 250 ng RNA (measured on a Nanodrop1000 spectrophotometer, ThermoScientific) were retro-transcribed using Maxima first strand cDNA synthesis kit (ThermoScientific) following the manufacturer’s instructions, including a dsDNAse treatment, in 20 µL final volume. For PCR, 0.5 µL cDNA were used as template for amplification using the Master Mix PCR (2X) kit (ThermoScientific) according to manufacturer’s instructions. The relative presence of the spliced and unspliced amplicons generated after RT-PCR was determined using a DNA12000 chip on a Bioanalyzer (Agilent) and analysed running the 2100expert software (v.B02.11.SI824). Splicing efficiency was calculated by comparing the area under the curves of electropherogram peaks corresponding to the different mRNA isoforms. Quantitative PCR was performed using QuantiNova SYBR Green PCR kit (Qiagen) according to the manufacturer’s instructions (Bustin et al. 2009). Primer sequence, RT-(q)PCR detailed conditions and amplicons extraction and purification are detailed in supplementary material.

#### Label-free proteomic analysis

Pelleted cells were lysed using RIPA buffer (50 mM Tris-HCl pH7.4, 150 mM NaCl, 1% Triton X-100, 0.5% Na deoxycholate, 1 mM EDTA) supplemented with 30 µg/mL of anti-protease mixture (Roche Diagnostics) during 20 min at 4°C. After centrifugation at 15,000x*g* for 10 min at 4°C, protein concentrations were determined in supernatant using Bio-Rad Protein Assay (Bio-Rad) according to the manufacturer’s instructions using Bovine Serum Albumin (Sigma) as standard. Label-free proteomic was performed at the Montpellier Proteomics Platform (PPM, BioCampus Montpellier), ran on all conditions and from three biological replicates. 20 µg of proteins were in-gel digested and resulting peptides were analysed using a Q Exactive HF mass spectrometer coupled with an Ultimate 3000 RSLC system (Thermo Fisher Scientific). MS/MS analyses were performed running the Maxquant software (v1.5.5.1). All MS/MS spectra were searched by the Andromeda search engine against a decoy database consisting in a combination of Homo sapiens entries from Reference Proteome (UP000005640, release 2019_02, https://www.uniprot.org/), a database with classical contaminants, and the sequences of interest (SHBLE, SHBLE*I, SHBLE*II and GFP). After excluding the usual contaminants, we obtained a final set of 4199 proteins detected at least once in one of the samples. Intensity based absolute quantification (iBAQ) normalised by the total iBAQ amount of each sample was used to compare heterologous protein levels between samples and replicates. Mass spectrometry proteomics data have been deposited to the ProteomeXchange Consortium via the PRIDE (Perez-Riverol et al. 2022) partner repository with the dataset identifier PXD047576 and 10.6019/PXD047576.

#### Data treatment

Data analyses and statistical analyses were carried out using R (v4.3.1) and R studio software (v 2023.06.1 Build 524). All packages used are included in the scripts available on git-hub (see data availability paragraph).

## Data availability

Vector and insert sequences are available on GenBank (pcDNA3.1Shble1_eGFP : OR659018, pcDNA3.1Shble2_eGFP: OR659019, pcDNA3.1Shble3_eGFP: OR659020, pcDNA3.1Shble4_eGFP: OR659021, pcDNA3.1Shble5_eGFP: OR659022, pcDNA3.1Shble6_eGFP: OR659023, pcDNA3.1Shble7_eGFP: OR659024, pcDNA3.1Shble8_eGFP: OR659025, pcDNA3.1Shble9_eGFP: OR659026, pcDNA3.1Shble10_eGFP: OR659027, pcDNA3.1Shble11_eGFP: OR659028, pcDNA3.1Shble12_eGFP: OR659029 and pcDNA3.1Shble13_eGFP: OR659030 and pcDNA3.1Shble4mut_eGFP: OR659031, pcDNA3.1Shble6mut_eGFP: OR659032, pcDNA3.1Shble7mut_eGFP: OR659033, pcDNA3.1Shble10mut_eGFP: OR659034 and pcDNA3.1Shble13mut_eGFP: OR659035). All R scripts used to analyse the data are available at <u>Github.com/philippe-paget/PERVASIVE-TRANSLATION-OF-THE-DOWNSTREAM-ORF-FROM-BICISTRONIC-MRNAS-BY-HUMAN-CELLS</u>. The input data .csv files used are also available along the scripts used for cytometry raw data. Raw data from cytometer, RTqPCR, Bioanalyzer and Sanger sequencing can be shared upon reasonable request to the corresponding author. Label-free proteomic data are available on the ProteomeXchange platform under the identifier PXD047576.

## Acknowledgements

We acknowledge the Montpellier Proteomics Platform (PPM, BioCampus Montpellier) for mass spectrometry experiments and the MRI imaging facility, member of the France-BioImaging national infrastructure supported by the French National Research Agency (ANR-10-INBS-04, «Investments for the future») for flow cytometry experiments.

## Funding and additional information

This study is supported by the European Union’s Horizon 2020 research and innovation program under the grant agreement CODOVIREVOL (ERC-2014-CoG-647916) to IGB. PPB is the recipient of a two-year post-doctoral grant from Fondation pour la Recherche Médicale.

## Supporting Information

Table S1: Composition variables of each construct, unspliced and spliced mRNAs.

Table S2: Primer sequences used

Figure S1: Vector map, mRNA produced and their general features. (panel A)

Figure S2: *shble* splicing events characterization.

Figure S3: Alternative CUPrefs of *shble* versions does not affect mRNA levels

Figure S4: Relation between *shble* translation efficiency and individual composition variables.

Figure S5: Determination of egfp translation efficiency.

Figure S6: SHBLE and GFP production from bicistronic *shble_egfp* mRNA.

Figure S7: Splicing efficiency determination.

Figure S8: Splicing does not affect mRNA levels.

Figure S9: Splicing effect on SHBLE and GFP translation efficiency.

## REFERENCES

Alexaki, Aikaterini, Jacob Kames, David D. Holcomb, John Athey, Luis V. Santana-Quintero, Phuc Vihn Nguyen Lam, Nobuko Hamasaki-Katagiri, et al. 2019. «Codon and Codon-Pair Usage Tables (CoCoPUTs): Facilitating Genetic Variation Analyses and Recombinant Gene Design». Journal of Molecular Biology 431 (13): 2434-41. 10.1016/j.jmb.2019.04.021.

Alonso, Andres M, et Luis Diambra. 2023. «Dicodon-Based Measures for Modeling Gene Expression. Édité par Tobias Marschall». Bioinformatics 39 (6): btad380. 10.1093/bioinformatics/btad380.

Arike, L., K. Valgepea, L. Peil, R. Nahku, K. Adamberg, et R. Vilu. 2012. «Comparison and Applications of Label-Free Absolute Proteome Quantification Methods on Escherichia Coli». Journal of Proteomics 75 (17): 5437-48. 10.1016/j.jprot.2012.06.020.

Bauer, Asli Petra, Doris Leikam, Simone Krinner, Frank Notka, Christine Ludwig, Gernot Längst, et Ralf Wagner. 2010. «The Impact of Intragenic CpG Content on Gene Expression». Nucleic Acids Research 38 (12): 3891-3908. 10.1093/nar/gkq115.

Bénitière, Florian, Anamaria Necsulea, et Laurent Duret. 2024. «Random Genetic Drift Sets an Upper Limit on mRNA Splicing Accuracy in Metazoans». eLife 13 (mars): RP93629. 10.7554/eLife.93629.

Beutler, E, T Gelbart, J H Han, J A Koziol, et B Beutler. 1989. «Evolution of the Genome and the Genetic Code: Selection at the Dinucleotide Level by Methylation and Polyribonucleotide Cleavage». Proceedings of the National Academy of Sciences 86 (1): 192–96. 10.1073/pnas.86.1.192.

Boël, Grégory, Reka Letso, Helen Neely, W. Nicholson Price, Kam-Ho Wong, Min Su, Jon D. Luff, et al. 2016. «Codon Influence on Protein Expression in E. Coli Correlates with mRNA Levels». Nature 529 (7586): 358–63. 10.1038/nature16509.

Bourret, Jérôme, Samuel Alizon, et Ignacio G Bravo. 2019. «COUSIN (COdon Usage Similarity INdex): A Normalized Measure of Codon Usage Preferences». Genome Biology and Evolution 11 (12): 3523–28. 10.1093/gbe/evz262.

Brinster, R L, J M Allen, R R Behringer, R E Gelinas, et R D Palmiter. 1988. «Introns Increase Transcriptional Efficiency in Transgenic Mice». Proceedings of the National Academy of Sciences 85 (3): 836–40. 10.1073/pnas.85.3.836.

Broadbent, Andrew J., Celia P. Santos, Amanda Anafu, Eckard Wimmer, Steffen Mueller, et Kanta Subbarao. 2016. «Evaluation of the Attenuation, Immunogenicity, and Efficacy of a Live Virus Vaccine Generated by Codon-Pair Bias de-Optimization of the 2009 Pandemic H1N1 Influenza Virus, in Ferrets». Vaccine 34 (4): 563-70. 10.1016/j.vaccine.2015.11.054.

Bustin, Stephen A, Vladimir Benes, Jeremy A Garson, Jan Hellemans, Jim Huggett, Mikael Kubista, Reinhold Mueller, et al. 2009. «The MIQE Guidelines: Minimum Information for Publication of Quantitative Real-Time PCR Experiments». Clinical Chemistry 55 (4): 611–22. 10.1373/clinchem.2008.112797.

Callens, Martijn, Léa Pradier, Michael Finnegan, Caroline Rose, et Stéphanie Bedhomme. 2021. «Read between the Lines: Diversity of Nontranslational Selection Pressures on Local Codon Usage. Édité par Paul Sharp». Genome Biology and Evolution 13 (9): evab097. 10.1093/gbe/evab097.

Callis, J., M. Fromm, et V. Walbot. 1987. «Introns Increase Gene Expression in Cultured Maize Cells». Genes & Development 1 (10): 1183–1200. 10.1101/gad.1.10.1183.

Calvo, Sarah E., David J. Pagliarini, et Vamsi K. Mootha. 2009. «Upstream Open Reading Frames Cause Widespread Reduction of Protein Expression and Are Polymorphic among Humans». Proceedings of the National Academy of Sciences 106 (18): 7507–12. 10.1073/pnas.0810916106.

Cambray, Guillaume, Joao C Guimaraes, et Adam Paul Arkin. 2018. «Evaluation of 244,000 Synthetic Sequences Reveals Design Principles to Optimize Translation in Escherichia Coli». Nature Biotechnology 36 (10): 1005–15. 10.1038/nbt.4238.

Chen, Jin, Andreas-David Brunner, J. Zachery Cogan, James K. Nuñez, Alexander P. Fields, Britt Adamson, Daniel N. Itzhak, et al. 2020. «Pervasive Functional Translation of Noncanonical Human Open Reading Frames». Science 367 (6482): 1140–46. 10.1126/science.aay0262.

Chen, Siyu, Ke Li, Wenqing Cao, Jia Wang, Tong Zhao, Qing Huan, Yu-Fei Yang, Shaohuan Wu, et Wenfeng Qian. 2017. «Codon-Resolution Analysis Reveals a Direct and Context-Dependent Impact of Individual Synonymous Mutations on mRNA Level». Molecular Biology and Evolution 34 (11): 2944–58. 10.1093/molbev/msx229.

Cheng, Jun, Kerstin C. Maier, Žiga Avsec, Petra Rus, et Julien Gagneur. 2017. «*Cis* - Regulatory Elements Explain Most of the mRNA Stability Variation across Genes in Yeast». RNA 23 (11): 1648–59. 10.1261/rna.062224.117.

Chew, Guo-Liang, Andrea Pauli, et Alexander F. Schier. 2016. «Conservation of uORF Repressiveness and Sequence Features in Mouse, Human and Zebrafish». Nature Communications 7 (1): 11663. 10.1038/ncomms11663.

Chin, Ju Xin, Bevan Kai-Sheng Chung, et Dong-Yup Lee. 2014. «Codon Optimization OnLine (COOL): A Web-Based Multi-Objective Optimization Platform for Synthetic Gene Design». Bioinformatics 30 (15): 2210–12. 10.1093/bioinformatics/btu192.

Chu, Dominique, Eleanna Kazana, Noémie Bellanger, Tarun Singh, Mick F Tuite, et Tobias Von Der Haar. 2014. «Translation Elongation Can Control Translation Initiation on Eukaryotic mRNAs». The EMBO Journal 33 (1): 21–34. 10.1002/embj.201385651.

Cigan, A. Mark, Lan Feng, et Thomas F. Donahue. 1988. «tRNA i ^met^ Functions in Directing the Scanning Ribosome to the Start Site of Translation». Science 242 (4875): 93–97. 10.1126/science.3051379.

Conrad, Steven J., Robert F. Silva, Cari J. Hearn, Megan Climans, et John R. Dunn. 2018. «Attenuation of Marek’s Disease Virus by Codon Pair Deoptimization of a Core Gene». Virology 516 (mars): 219–26. 10.1016/j.virol.2018.01.020.

Courel, Maïté, Yves Clément, Clémentine Bossevain, Dominika Foretek, Olivia Vidal Cruchez, Zhou Yi, Marianne Bénard, et al. 2019. «GC Content Shapes mRNA Storage and Decay in Human Cells». eLife 8 (décembre): e49708. 10.7554/eLife.49708.

Daron, Josquin, et Ignacio Bravo. 2021. «Variability in Codon Usage in Coronaviruses Is Mainly Driven by Mutational Bias and Selective Constraints on CpG Dinucleotide». Viruses 13 (9): 1800. 10.3390/v13091800.

Desmet, François-Olivier, Dalil Hamroun, Marine Lalande, Gwenaëlle Collod-Béroud, Mireille Claustres, et Christophe Béroud. 2009. «Human Splicing Finder: an online bioinformatics tool to predict splicing signals». Nucleic Acids Research 37 (9): e67. 10.1093/nar/gkp215.

Doyle, Francis, Andrea Leonardi, Lauren Endres, Scott A. Tenenbaum, Peter C. Dedon, et Thomas J. Begley. 2016. «Gene-and Genome-Based Analysis of Significant Codon Patterns in Yeast, Rat and Mice Genomes with the CUT Codon UTilization Tool». Methods 107 (septembre): 98–109. 10.1016/j.ymeth.2016.05.010.

Duan, Jubao, et Marcos A. Antezana. 2003. «Mammalian Mutation Pressure, Synonymous Codon Choice, and mRNA Degradation». Journal of Molecular Evolution 57 (6): 694–701. 10.1007/s00239-003-2519-1.

Eldemery, Fatma, Changbo Ou, Taejoong Kim, Stephen Spatz, John Dunn, Robert Silva, et Qingzhong Yu. 2023. «Evaluation of Newcastle Disease Virus LaSota Strain Attenuated by Codon Pair Deoptimization of the HN and F Genes for in Ovo Vaccination». Veterinary Microbiology 277 (février): 109625. 10.1016/j.vetmic.2022.109625.

Eskesen, S T, F N Eskesen, et A Ruvinsky. 2004. «Natural Selection Affects Frequencies of AG and GT Dinucleotides at the 5′ and 3′ Ends of Exons». Genetics 167 (1): 543–50. 10.1534/genetics.167.1.543.

Fallot, Stéphanie, Raouia Ben Naya, Corinne Hieblot, Philippe Mondon, Eric Lacazette, Khalil Bouayadi, Abdelhakim Kharrat, Christian Touriol, et Hervé Prats. 2009. «Alternative-Splicing-Based Bicistronic Vectors for Ratio-Controlled Protein Expression and Application to Recombinant Antibody Production.» Nucleic Acids Research 37 (20): e134-e134. 10.1093/nar/gkp716.

Fu, Hongguang, Yanbing Liang, Xiuqin Zhong, ZhiLing Pan, Lei Huang, HaiLin Zhang, Yang Xu, Wei Zhou, et Zhong Liu. 2020. «Codon Optimization with Deep Learning to Enhance Protein Expression». Scientific Reports 10 (1): 17617. 10.1038/s41598-020-74091-z.

Gao, Feng, Yingying Li, Julie M. Decker, Fred W. Peyerl, Frederic Bibollet-Ruche, Cynthia M. Rodenburg, Yalu Chen, et al. 2003. «Codon Usage Optimization of HIV Type 1 Subtype C *Gag*, *Pol*, *Env*, and *Nef* Genes: *In Vitro* Expression and Immune Responses in DNA-Vaccinated Mice». AIDS Research and Human Retroviruses 19 (9): 817–23. 10.1089/088922203769232610.

Gao, Kaiping, Akio Masuda, Tohru Matsuura, et Kinji Ohno. 2008. «Human Branch Point Consensus Sequence Is yUnAy». Nucleic Acids Research 36 (7): 2257–67. 10.1093/nar/gkn073.

Graf, Marcus, Ludwig Deml, et Ralf Wagner. 2003. Codon-Optimized Genes That Enable Increased Heterologous Expression in Mammalian Cells and Elicit Efficient Immune Responses in Mice after Vaccination of Naked DNA. In Molecular Diagnosis of Infectious Diseases, par Jochen Decker et Udo Reischl, 0:197-210. New Jersey: Humana Press. 10.1385/1-59259-679-7:197.

Hanson, Gavin, et Jeff Coller. 2018. «Codon Optimality, Bias and Usage in Translation and mRNA Decay». Nature Reviews Molecular Cell Biology 19 (1): 20–30. 10.1038/nrm.2017.91.

Hellens, Roger P., Chris M. Brown, Matthew A.W. Chisnall, Peter M. Waterhouse, et Richard C. Macknight. 2016. «The Emerging World of Small ORFs». Trends in Plant Science 21 (4): 317–28. 10.1016/j.tplants.2015.11.005.

Hernandez-Alias, Xavier, Hannah Benisty, Leandro G. Radusky, Luis Serrano, et Martin H. Schaefer. 2023. «Using Protein-per-mRNA Differences among Human Tissues in Codon Optimization». Genome Biology 24 (1): 34. 10.1186/s13059-023-02868-2.

Hinnebusch, Alan G., Ivaylo P. Ivanov, et Nahum Sonenberg. 2016. «Translational Control by 5′-Untranslated Regions of Eukaryotic mRNAs». Science 352 (6292): 1413–16. 10.1126/science.aad9868.

Ingolia, Nicholas T., Liana F. Lareau, et Jonathan S. Weissman. 2011. «Ribosome Profiling of Mouse Embryonic Stem Cells Reveals the Complexity and Dynamics of Mammalian Proteomes». Cell 147 (4): 789–802. 10.1016/j.cell.2011.10.002.

Kaishima, Misato, Jun Ishii, Toshihide Matsuno, Nobuo Fukuda, et Akihiko Kondo. 2016. «Expression of Varied GFPs in Saccharomyces Cerevisiae: Codon Optimization Yields Stronger than Expected Expression and Fluorescence Intensity». Scientific Reports 6 (1): 35932. 10.1038/srep35932.

Kaufman, R.J., P. Murtha, et M.V. Davies. 1987. «Translational Efficiency of Polycistronic mRNAs and Their Utilization to Express Heterologous Genes in Mammalian Cells». The EMBO Journal 6 (1): 187–93. 10.1002/j.1460-2075.1987.tb04737.x.

Kosovac, D, J Wild, C Ludwig, S Meissner, A P Bauer, et R Wagner. 2011. «Minimal Doses of a Sequence-Optimized Transgene Mediate High-Level and Long-Term EPO Expression in Vivo: Challenging CpG-Free Gene Design». Gene Therapy 18 (2): 189–98. 10.1038/gt.2010.134.

Kozak, M. 1986. «Influences of mRNA Secondary Structure on Initiation by Eukaryotic Ribosomes». Proceedings of the National Academy of Sciences 83 (9): 2850–54. 10.1073/pnas.83.9.2850.

Kozak, M. 1989. «The Scanning Model for Translation: An Update». The Journal of Cell Biology 108 (2): 229–41. 10.1083/jcb.108.2.229.

Kudla, Grzegorz, Andrew W. Murray, David Tollervey, et Joshua B. Plotkin. 2009. «Coding-Sequence Determinants of Gene Expression in *Escherichia Coli*». Science 324 (5924): 255–58. 10.1126/science.1170160.

Kunec, Dusan, et Nikolaus Osterrieder. 2016. «Codon Pair Bias Is a Direct Consequence of Dinucleotide Bias». Cell Reports 14 (1): 55–67. 10.1016/j.celrep.2015.12.011.

Kwon, Kwang-Chul, Hui-Ting Chan, Ileana R. León, Rosalind Williams-Carrier, Alice Barkan, et Henry Daniell. 2016. «Codon Optimization to Enhance Expression Yields Insights into Chloroplast Translation». Plant Physiology 172 (1): 62–77. 10.1104/pp.16.00981.

Le Hir, Hervé, Ajit Nott, et Melissa J. Moore. 2003. «How Introns Influence and Enhance Eukaryotic Gene Expression». Trends in Biochemical Sciences 28 (4): 215–20. 10.1016/S0968-0004(03)00052-5.

Lim, Chun Shen, Samuel J T. Wardell, Torsten Kleffmann, et Chris M Brown. 2018. «The Exon–Intron Gene Structure Upstream of the Initiation Codon Predicts Translation Efficiency». Nucleic Acids Research 46 (9): 4575–91. 10.1093/nar/gky282.

Lin, Yizhu, Gemma E May, Hunter Kready, Lauren Nazzaro, Mao Mao, Pieter Spealman, Yehuda Creeger, et C Joel McManus. 2019. «Impacts of uORF Codon Identity and Position on Translation Regulation». Nucleic Acids Research 47 (17): 9358–67. 10.1093/nar/gkz681.

Liu, Ziqing, Olivia Chen, J. Blake Joseph Wall, Michael Zheng, Yang Zhou, Li Wang, Haley Ruth Vaseghi, Li Qian, et Jiandong Liu. 2017. «Systematic Comparison of 2A Peptides for Cloning Multi-Genes in a Polycistronic Vector». Scientific Reports 7 (1): 2193. 10.1038/s41598-017-02460-2.

Luo, M. J., et R. Reed. 1999. «Splicing Is Required for Rapid and Efficient mRNA Export in Metazoans». Proceedings of the National Academy of Sciences of the United States of America 96 (26): 14937–42. 10.1073/pnas.96.26.14937.

Lynch, Michael, et Georgi K. Marinov. 2015. «The Bioenergetic Costs of a Gene». Proceedings of the National Academy of Sciences 112 (51): 15690–95. 10.1073/pnas.1514974112.

Lyu, Xueliang, Qian Yang, Fangzhou Zhao, et Yi Liu. 2021. «Codon Usage and Protein Length-Dependent Feedback from Translation Elongation Regulates Translation Initiation and Elongation Speed». Nucleic Acids Research 49 (16): 9404–23. 10.1093/nar/gkab729.

Markham, Nicholas R., et Michael Zuker. 2008. UNAFold. In Bioinformatics: Structure, Function and Applications, édité par Jonathan M. Keith, 3-31. Methods in Molecular Biology^TM^. Totowa, NJ: Humana Press. 10.1007/978-1-60327-429-6_1.

Matsumoto, K., K. M. Wassarman, et A. P. Wolffe. 1998. «Nuclear History of a Pre-mRNA Determines the Translational Activity of Cytoplasmic mRNA». The EMBO Journal 17 (7): 2107–21. 10.1093/emboj/17.7.2107.

Mauro, Vincent P., et Stephen A. Chappell. 2014. «A Critical Analysis of Codon Optimization in Human Therapeutics». Trends in Molecular Medicine 20 (11): 604–13. 10.1016/j.molmed.2014.09.003.

Mertz, J E, A Murphy, et A Barkan. 1983. «Mutants Deleted in the Agnogene of Simian Virus 40 Define a New Complementation Group». Journal of Virology 45 (1): 36–46. 10.1128/jvi.45.1.36-46.1983.

Mordstein, Christine, Rosina Savisaar, Robert S. Young, Jeanne Bazile, Lana Talmane, Juliet Luft, Michael Liss, Martin S. Taylor, Laurence D. Hurst, and Grzegorz Kudla. 2020. «Codon Usage and Splicing Jointly Influence mRNA Localization». Cell Systems 10 (4): 351–362.e8. 10.1016/j.cels.2020.03.001.

Nagata, Toshi, Masato Uchijima, Atsushi Yoshida, Minae Kawashima, et Yukio Koide. 1999. «Codon Optimization Effect on Translational Efficiency of DNA Vaccine in Mammalian Cells: Analysis of Plasmid DNA Encoding a CTL Epitope Derived from Microorganisms». Biochemical and Biophysical Research Communications 261 (2): 445–51. 10.1006/bbrc.1999.1050.

Newman, Zachary R., Janet M. Young, Nicholas T. Ingolia, et Gregory M. Barton. 2016. «Differences in Codon Bias and GC Content Contribute to the Balanced Expression of TLR7 and TLR9». Proceedings of the National Academy of Sciences 113 (10). 10.1073/pnas.1518976113.

Nogales, Aitor, Steven F. Baker, Emilio Ortiz-Riaño, Stephen Dewhurst, David J. Topham, et Luis Martínez-Sobrido. 2014. «Influenza A Virus Attenuation by Codon Deoptimization of the NS Gene for Vaccine Development». Édité par T. S. Dermody. Journal of Virology 88 (18): 10525–40. 10.1128/JVI.01565-14.

Odon, Valerie, Jelke J Fros, Niluka Goonawardane, Isabelle Dietrich, Ahmad Ibrahim, Kinda Alshaikhahmed, Dung Nguyen, et Peter Simmonds. 2019. «The Role of ZAP and OAS3/RNAseL Pathways in the Attenuation of an RNA Virus with Elevated Frequencies of CpG and UpA Dinucleotides». Nucleic Acids Research 47 (15): 8061–83. 10.1093/nar/gkz581.

Peabody, D S, et P Berg. 1986. «Termination-Reinitiation Occurs in the Translation of Mammalian Cell mRNAs». Molecular and Cellular Biology 6 (7): 2695–2703. 10.1128/MCB.6.7.2695.

Peabody, David S., Suresh Subramani, et Paul Berg. 1986. «Effect of Upstream Reading Frames on Translation Efficiency in Simian Virus 40 Recombinants». Molecular and Cellular Biology 6 (7): 2704–11. 10.1128/mcb.6.7.2704-2711.1986.

Perez-Riverol, Yasset, Jingwen Bai, Chakradhar Bandla, David García-Seisdedos, Suresh Hewapathirana, Selvakumar Kamatchinathan, Deepti J Kundu, et al. 2022. «The PRIDE Database Resources in 2022: A Hub for Mass Spectrometry-Based Proteomics Evidences». Nucleic Acids Research 50 (D1): D543–52. 10.1093/nar/gkab1038.

Picard, Marion A. L., Fiona Leblay, Cécile Cassan, Anouk Willemsen, Josquin Daron, Frédérique Bauffe, Mathilde Decourcelle, Antonin Demange, et Ignacio G. Bravo. 2023. «Transcriptomic, Proteomic, and Functional Consequences of Codon Usage Bias in Human Cells during Heterologous Gene Expression». Protein Science 32 (3): e4576. 10.1002/pro.4576.

Pop, Cristina, Silvi Rouskin, Nicholas T Ingolia, Lu Han, Eric M Phizicky, Jonathan S Weissman, et Daphne Koller. 2014. «Causal Signals between Codon Bias, MRNA Structure, and the Efficiency of Translation and Elongation». Molecular Systems Biology 10 (12): 770. 10.15252/msb.20145524.

Powell, Michael L., T. David K. Brown, et Ian Brierley. 2008. «Translational Termination-Re-Initiation in Viral Systems». Biochemical Society Transactions 36 (Pt 4): 717–22. 10.1042/BST0360717.

Pöyry, Tuija A. A., Ann Kaminski, Emma J. Connell, Christopher S. Fraser, et Richard J. Jackson. 2007. «The Mechanism of an Exceptional Case of Reinitiation after Translation of a Long ORF Reveals Why Such Events Do Not Generally Occur in Mammalian mRNA Translation». Genes & Development 21 (23): 3149–62. 10.1101/gad.439507.

Presnyak, Vladimir, Najwa Alhusaini, Ying-Hsin Chen, Sophie Martin, Nathan Morris, Nicholas Kline, Sara Olson, et al. 2015. «Codon Optimality Is a Major Determinant of mRNA Stability». Cell 160 (6): 1111–24. 10.1016/j.cell.2015.02.029.

Puigbo, P., E. Guzman, A. Romeu, et S. Garcia-Vallve. 2007. «OPTIMIZER: A Web Server for Optimizing the Codon Usage of DNA Sequences». Nucleic Acids Research 35 (Web Server): W126-31. 10.1093/nar/gkm219.

Radrizzani, Sofia, Grzegorz Kudla, Zsuzsanna Izsvák, et Laurence D. Hurst. 2024. «Selection on Synonymous Sites: The Unwanted Transcript Hypothesis». Nature Reviews Genetics, janvier. 10.1038/s41576-023-00686-7.

Sandhu, Kuljeet Singh, Sunil Pandey, Souvik Maiti, et Beena Pillai. 2008. «GASCO: Genetic Algorithm Simulation for Codon Optimization». In Silico Biology 8 (2): 187–92.

Sharp, Paul M., et Wen-Hsiung Li. 1987. «The Codon Adaptation Index-a Measure of Directional Synonymous Codon Usage Bias, and Its Potential Applications». Nucleic Acids Research 15 (3): 1281–95. 10.1093/nar/15.3.1281.

Sibley, Christopher R., Lorea Blazquez, et Jernej Ule. 2016. «Lessons from Non-Canonical Splicing». Nature Reviews Genetics 17 (7): 407–21. 10.1038/nrg.2016.46.

Simmonds, Peter, Wenjun Xia, J Baillie, et Ken McKinnon. 2013. «Modelling Mutational and Selection Pressures on Dinucleotides in Eukaryotic Phyla –Selection against CpG and UpA in Cytoplasmically Expressed RNA and in RNA Viruses». BMC Genomics 14 (1): 610. 10.1186/1471-2164-14-610.

Solovyev, V. 2003. Statistical Approaches in Eukaryotic Gene Prediction. In Handbook of Statistical Genetics, édité par D. J. Balding, M. Bishop, et C. Cannings, 1^re^ éd. Wiley. 10.1002/0470022620.bbc06.

Sorokin, Ivan I., Konstantin S. Vassilenko, Ilya M. Terenin, Natalia O. Kalinina, Vadim I. Agol, et Sergey E. Dmitriev. 2021. «Non-Canonical Translation Initiation Mechanisms Employed by Eukaryotic Viral mRNAs». Biochemistry (Moscow) 86 (9): 1060–94. 10.1134/S0006297921090042.

Subramani, Suresh, Richard Mulligan, et Paul Berg. 1981. «Expression of the Mouse Dihydrofolate Reductase Complementary Deoxyribonucleic Acid in Simian Virus 40 Vectors». Molecular and Cellular Biology 1 (9): 854–64. 10.1128/mcb.1.9.854-864.1981.

Swartz, M.N., T.A. Trautner, et Arthur Kornberg. 1962. «Enzymatic Synthesis of Deoxyribonucleic Acid». Journal of Biological Chemistry 237 (6): 1961–67. 10.1016/S0021-9258(19)73967-2.

Takata, Matthew A., Daniel Gonçalves-Carneiro, Trinity M. Zang, Steven J. Soll, Ashley York, Daniel Blanco-Melo, et Paul D. Bieniasz. 2017. «CG Dinucleotide Suppression Enables Antiviral Defence Targeting Non-Self RNA». Nature 550 (7674): 124–27. 10.1038/nature24039.

Victor, Manish P., Debarun Acharya, Tina Begum, et Tapash C. Ghosh. 2019. «The Optimization of mRNA Expression Level by Its Intrinsic Properties—Insights from Codon Usage Pattern and Structural Stability of mRNA». Genomics 111 (6): 1292–97. 10.1016/j.ygeno.2018.08.009.

Wacholder, Aaron, et Anne-Ruxandra Carvunis. 2023. «Biological Factors and Statistical Limitations Prevent Detection of Most Noncanonical Proteins by Mass Spectrometry». PLoS Biology 21 (12): e3002409. 10.1371/journal.pbio.3002409.

Walsh, Derek, et Ian Mohr. 2011. «Viral Subversion of the Host Protein Synthesis Machinery». Nature Reviews Microbiology 9 (12): 860–75. 10.1038/nrmicro2655.

Wang, Zefeng, et Christopher B. Burge. 2008. «Splicing Regulation: From a Parts List of Regulatory Elements to an Integrated Splicing Code». RNA 14 (5): 802–13. 10.1261/rna.876308.

Yang, Qian, Xueliang Lyu, Fangzhou Zhao, et Yi Liu. 2021. «Effects of Codon Usage on Gene Expression Are Promoter Context Dependent». Nucleic Acids Research 49 (2): 818–31. 10.1093/nar/gkaa1253.

Yu, Chien-Hung, Yunkun Dang, Zhipeng Zhou, Cheng Wu, Fangzhou Zhao, Matthew S. Sachs, et Yi Liu. 2015. «Codon Usage Influences the Local Rate of Translation Elongation to Regulate Co-Translational Protein Folding». Molecular Cell 59 (5): 744–54. 10.1016/j.molcel.2015.07.018.

Zhao, Fangzhou, Chien-hung Yu, et Yi Liu. 2017. «Codon Usage Regulates Protein Structure and Function by Affecting Translation Elongation Speed in Drosophila Cells». Nucleic Acids Research 45 (14): 8484–92. 10.1093/nar/gkx501.

Zhao, Fangzhou, Zhipeng Zhou, Yunkun Dang, Hyunsoo Na, Catherine Adam, Anna Lipzen, Vivian Ng, Igor V. Grigoriev, et Yi Liu. 2021. «Genome-Wide Role of Codon Usage on Transcription and Identification of Potential Regulators». Proceedings of the National Academy of Sciences 118 (6): e2022590118. 10.1073/pnas.2022590118.

Zhao, Lei, Xiaoji Cong, Linhui Zhai, Hao Hu, Jun-Yu Xu, Wensi Zhao, Mengdi Zhu, Minjia Tan, et Bang-Ce Ye. 2020. «Comparative Evaluation of Label-Free Quantification Strategies». Journal of Proteomics 215 (mars): 103669. 10.1016/j.jprot.2020.103669.

Zhou, Zhipeng, Yunkun Dang, Mian Zhou, Lin Li, Chien-hung Yu, Jingjing Fu, She Chen, et Yi Liu. 2016. «Codon Usage Is an Important Determinant of Gene Expression Levels Largely through Its Effects on Transcription». Proceedings of the National Academy of Sciences 113 (41). 10.1073/pnas.1606724113.

