## Supplementary files for "Pervasive translation of the downstream ORF from bicistronic mRNAs by human cells: impact of the upstream ORF codon usage and splicing"

### Supporting information

**Table S1: Composition variables of each constructs, unspliced and spliced mRNAs.**
**Table S2: (separate .csv file) Data from label free proteomic deposited on the PRIDE database.**
**Table S3: Primer sequences used**

**Figure S1: Vector map, mRNA produced and their general features.**
**Figure S2 : *shble* splicing events characterization.**
**Figure S3: Alternative CUPrefs of *shble* versions does not affect mRNA levels.**
**Figure S4: Relation between *shble* translation efficiency and individual composition variables.**
**Figure S5: Determination of egfp translation efficiency.**
**Figure S6: SHBLE and GFP production from bicistronic *shble_egfp* mRNA.**
**Figure S7: Splicing efficiency determination.**
**Figure S8: Splicing does not affect mRNA levels.**
**Figure S9: Splicing effect on SHBLE and GFP translation efficiency.**

A

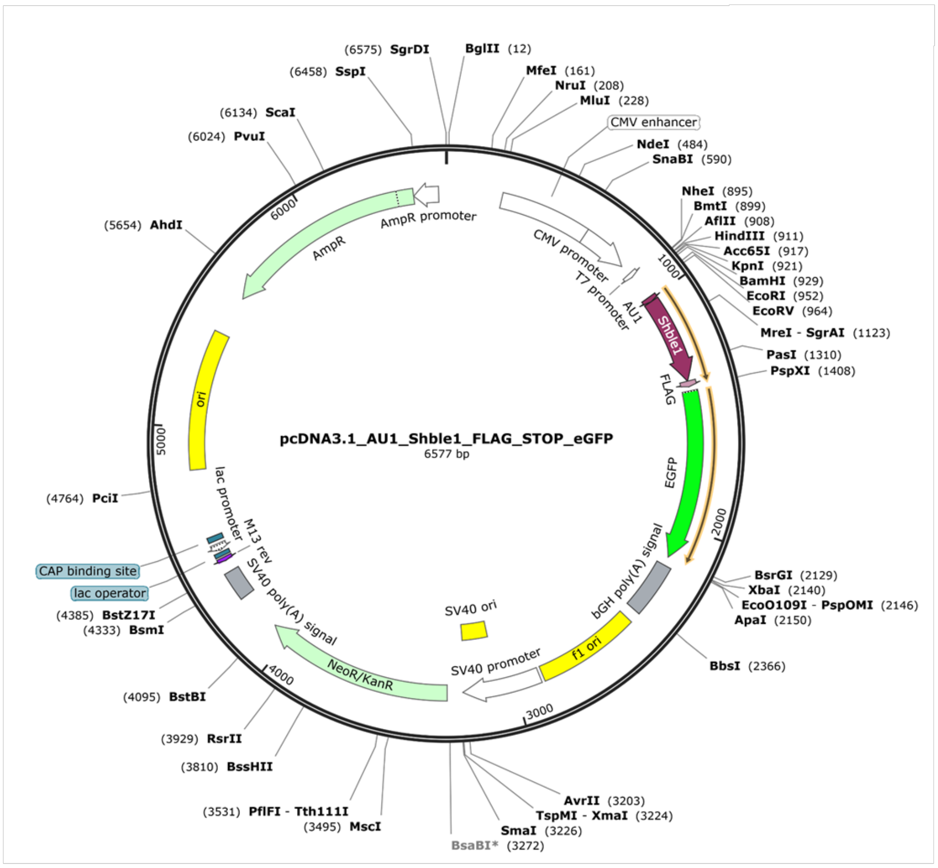


B

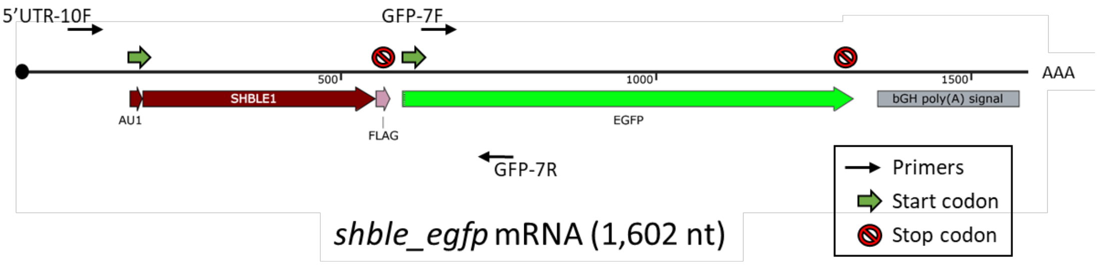


**Figure S1: Vector map, mRNA produced and their general features. (panel A)** Vector map showing the organization of the *shble* and *egfp* tandem ORF. Information available on GenBank (see data availability paragraph). **(panel B)** Organisation of the prototypic unspliced bicistronic *shble_egfp* mRNA. Are displayed ORF and their respective translation start and stop codons and the position of primers used for RT-PCR and RT-qPCR.

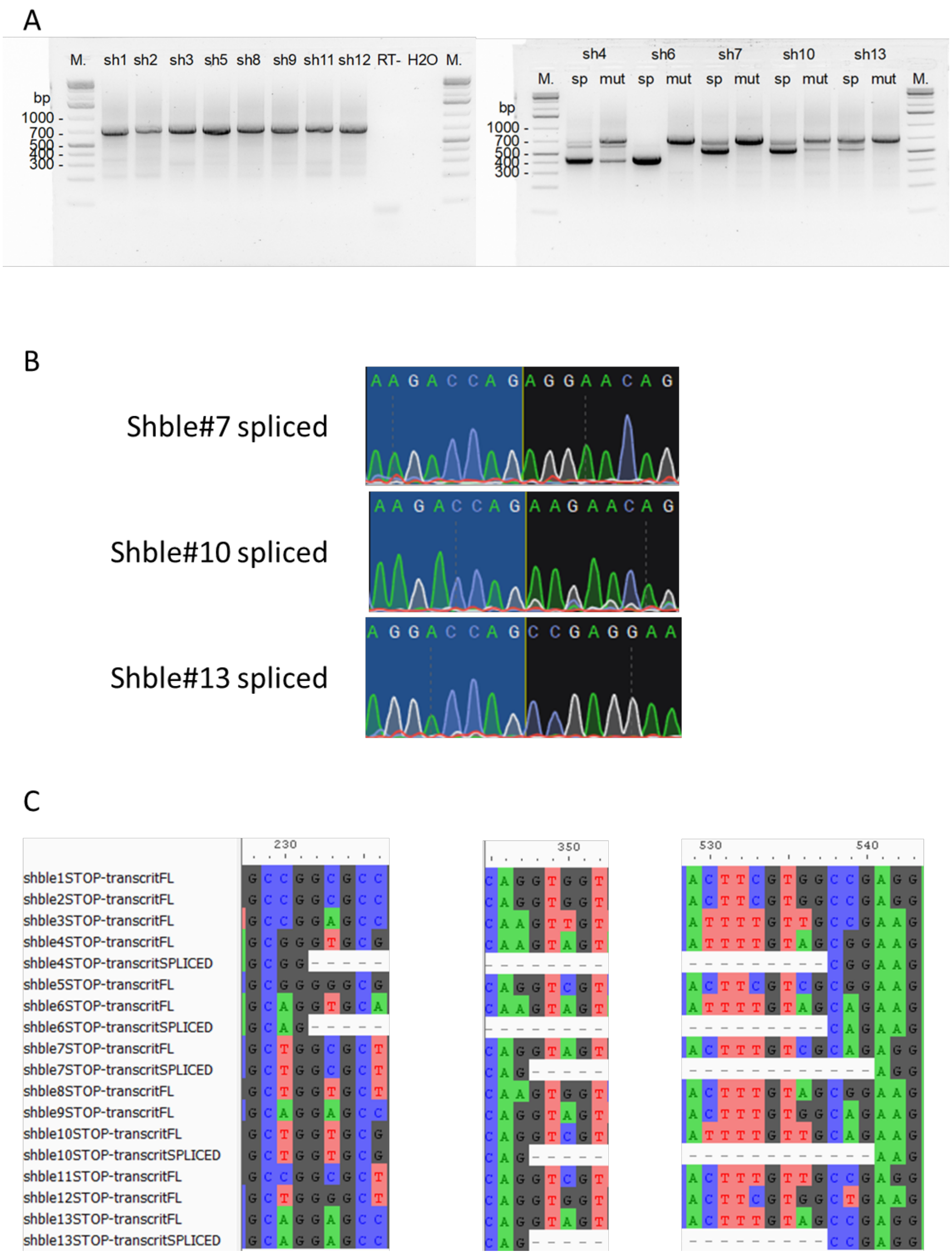


**Figure S2 : *shble* splicing events characterization. (panel A)** Agarose gel showing *shble_egfp* mRNA amplification by RT-PCR. M., molecular weight marker; RT-, negative control without reverse transcriptase; H2O, PCR without template. Figure should be read as follows, using *shble*#4 as an example: “sh4sp” refers to the sequence that undergoes splicing, while “sh4mut” refers to the sequence that has been mutated to ablate splicing. **(panel B)** Electropherograms of Sanger sequencing coming from the newly discovered and isolated spliced mRNA from *shble*#7, *shble*#10 and *shble*#13 showing splice junctions (blue=exon1, black=exon2). Bands were isolated and purified from an agarose gel after RT-PCR. **(panel C)** Alignment of the mRNAs isoform sequences produced by the thirteen *shble* constructs. Top number indicate position on the mRNA from the transcription start site. For spliced mRNA, dashed position correspond to the excised introns. All sequences are available on Genbank (see data availability paragraph).

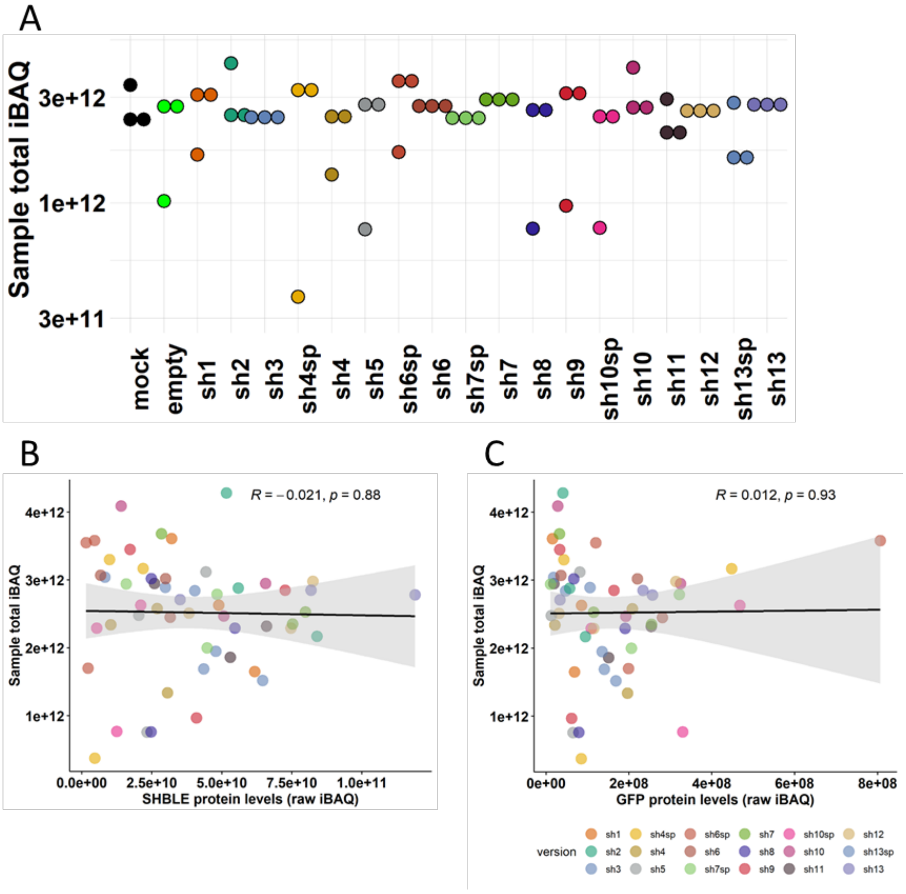


**Figure S3: Alternative CUPrefs of *shble* versions does not affect mRNA levels. (Panel A)** Dot-plot showing total iBAQ values for the three biological replicates used in figure 3. **(panel B and C)** Pearson's linear regression (black line) and 95% confidence interval of the fit (grey) between SHBLE or GFP iBAQ values, and total sample iBAQ values. Figure should be read as follows, using *shble*#4 as an example: "sh4sp" refers to the sequence that undergoes splicing, while "sh4" refers to the sequence that has been mutated to ablate splicing.

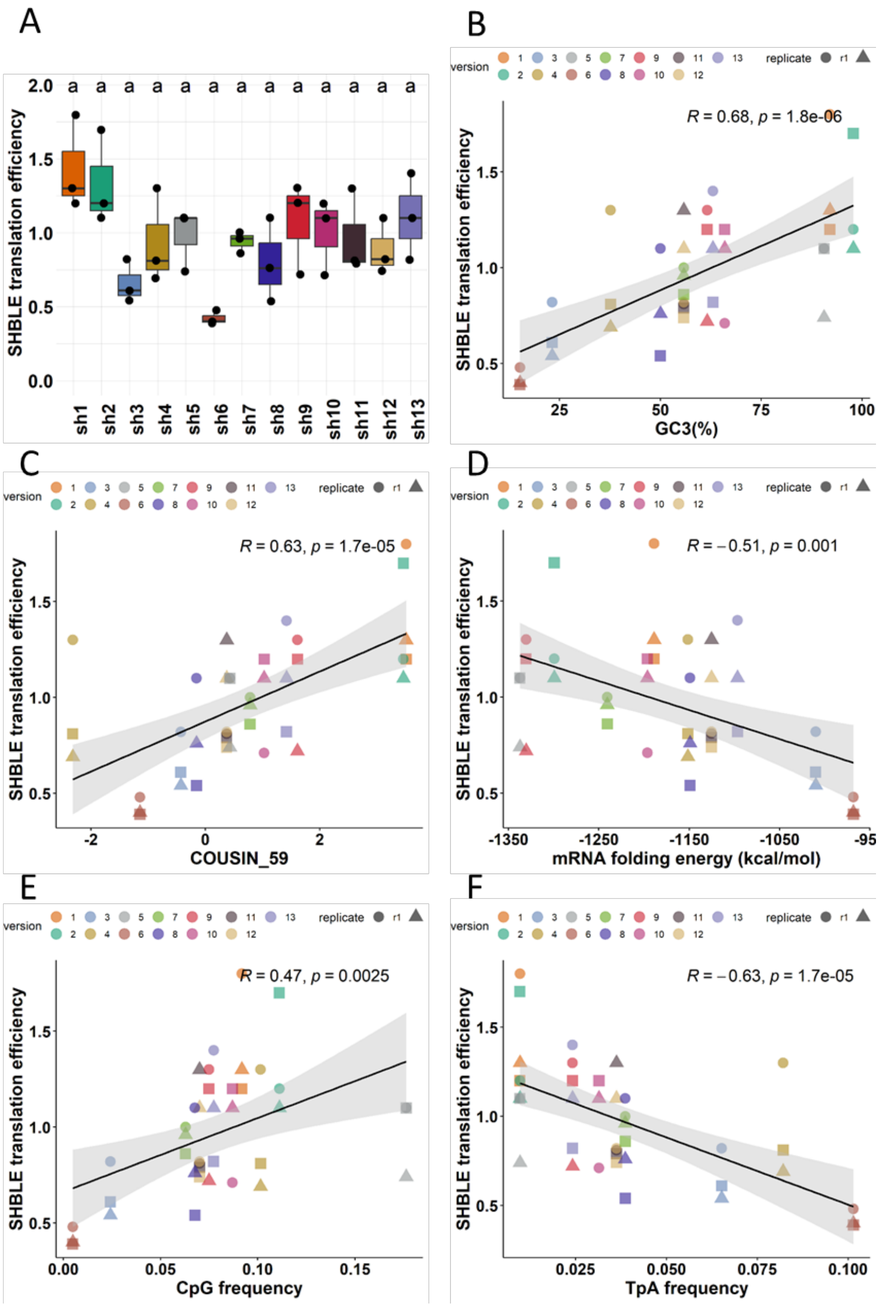


**Figure S4: Relation between *shble* translation efficiency and individual composition variables. (panel A)** Box-and-whiskers plot showing *shble* translation efficiency from the thirteen *shble* versions. Relative mRNA levels values and SHBLE protein levels values were normalised by the median value of the samples in the same replicate. Translation efficiency is calculated as protein-over-mRNA levels. Letters present the results of a pairwise Wilcoxon rank sum test with B-H adjusted p-values; $\alpha$=0.05; median values for samples labelled with the same letter are not statistically different. (**panels B to F**) Pearson's linear regression (black line) and 95% confidence interval of the fit (grey) between SHBLE translation efficiency, and **(panel B)** GC3, **(panel C)** COUSIN_59 score, **(panel D)** mRNA folding energy, **(panel E)** CpG dinucleotide frequency and **(panel F)** TpA dinucleotide frequency. For all panels, values from a same biological replicate are represented by triangle, rectangle or circle shapes.

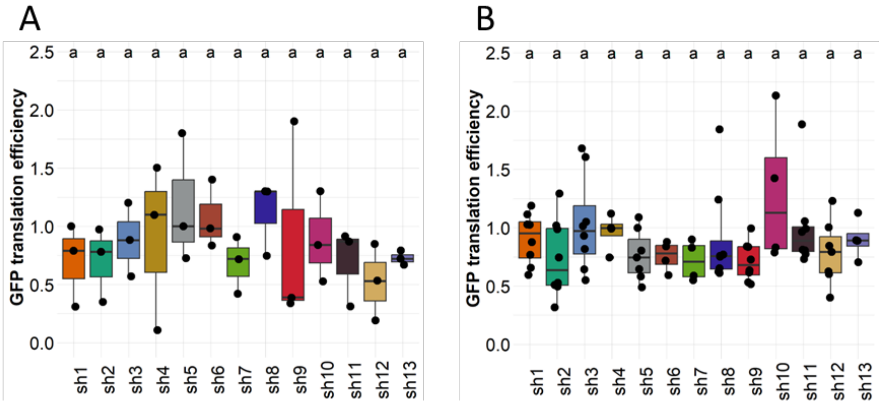


**Figure S5: Determination of egfp translation efficiency.** Box-and-whiskers plot showing *egfp* translation efficiency from the thirteen *shble* versions. Translation efficiency was determined by either normalising label free proteomic GFP protein levels **(panel A)** or cytometry GFP sums of fluorescence **(panel B)** to the relative mRNA levels. Relative mRNA levels values, GFP protein levels values or GFP sums on fluorescence were normalised by the median value of the samples in the same replicate. Letters present the results of a pairwise Wilcoxon rank sum test with B-H adjusted p-values; $\alpha$=0.05; median values for samples labelled with the same letter are not statistically different.

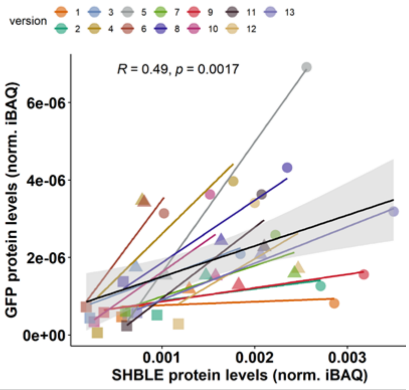


**Figure S6: SHBLE and GFP production from bicistronic *shble_egfp* mRNA.** Pearson's linear regression (coloured lines) between normalised SHBLE iBAQ levels and normalised GFP iBAQ levels from three biological replicates for each of the thirteen *shble* versions. The global Pearson linear regression (black line) and 95% confidence interval of the fit (grey) represent the combined fit of the thirteen versions. Values from a same biological replicate are represented by triangle, rectangle or circle shapes.

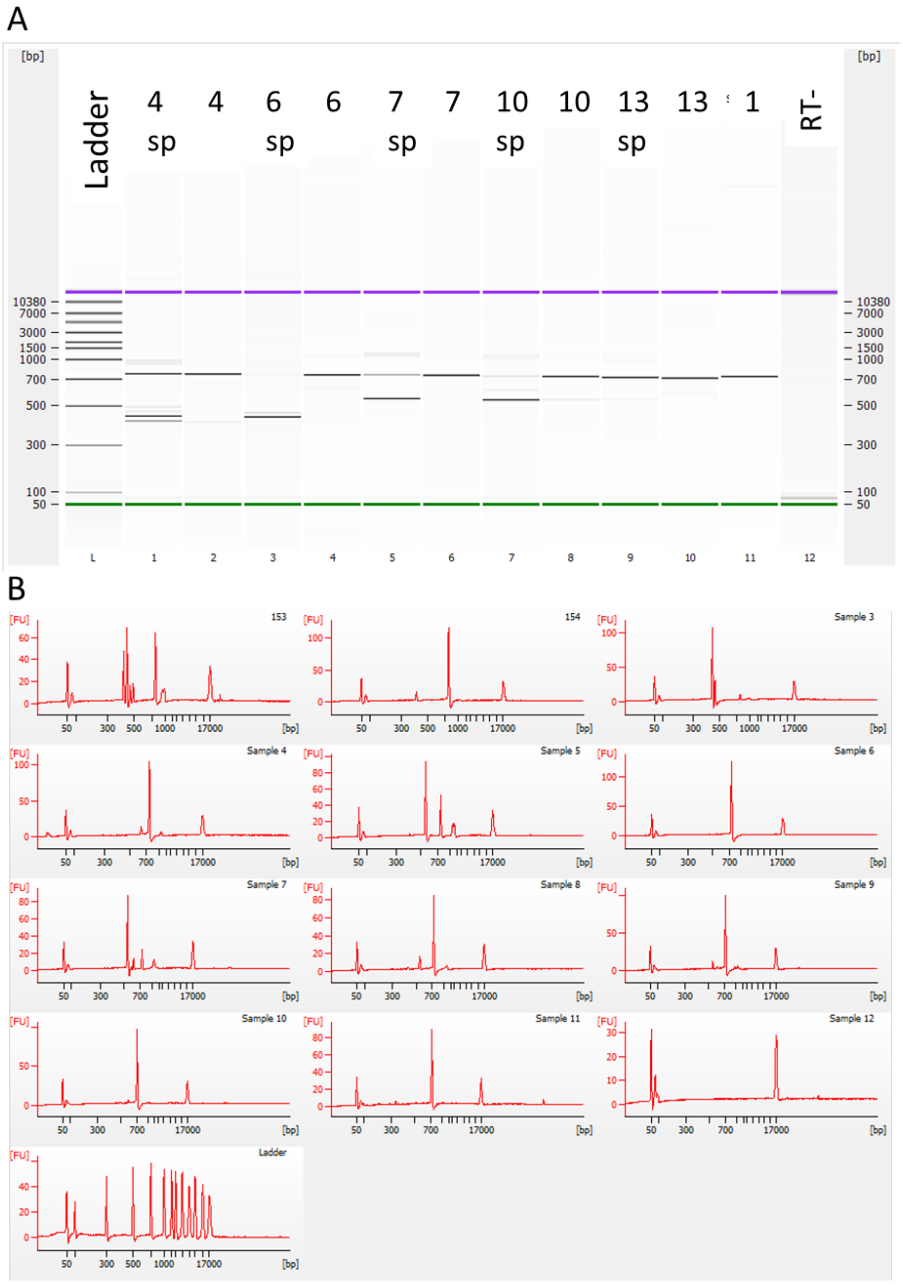


**Figure S7: Splicing efficiency determination. (panel A)** Gel migration on Bioanalyzer ship of RT-PCR amplicon targeting bicistronic *shble-egfp* mRNA coming from conditions transfected with constructs *shble*#4, #6, #7, #10 and #13, their splice-ablated counterparts, *shble*#1 as positive control and an RT- negative control**. (panel B)** Electropherogram of the same migration showing the fluorescent intensity of amplicons corresponding to spliced or unspliced bicistronic mRNAs. Bands between 400 and 500 bp correspond to the two splice products and band at 700 bp correspond to the unspliced mRNA. Bands at 50 and 17000 bp correspond to the control ladder.

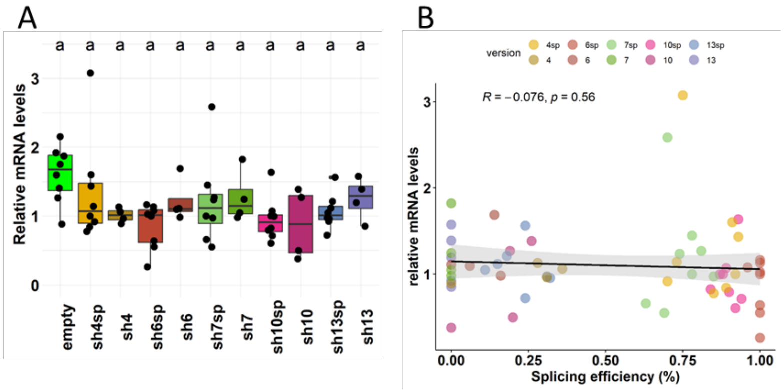


**Figure S8: Splicing does not affect mRNA levels. (panel A)** Box-and-whiskers plot showing relative levels of total heterologous mRNAs measured by RT-qPCR from four to eight biological replicates. For each replicates, relative mRNA levels values were normalised by the median value of samples. The positive control "empty" condition expressed monocistronic *egfp* coding mRNAs while the ten *shble* conditions expressed spliced and unspliced bicistronic *shble-egfp* mRNAs. (pairwise Wilcoxon rank sum test with B-H adjusted p-values ; α=0.05 ; Same letter mean no significant statistical difference). **(panel B)** Pearson's linear regression (black line) and 95% confidence interval of the fit (grey) between the splicing efficiency and the relative mRNA levels expressed in the ten *shble* conditions.

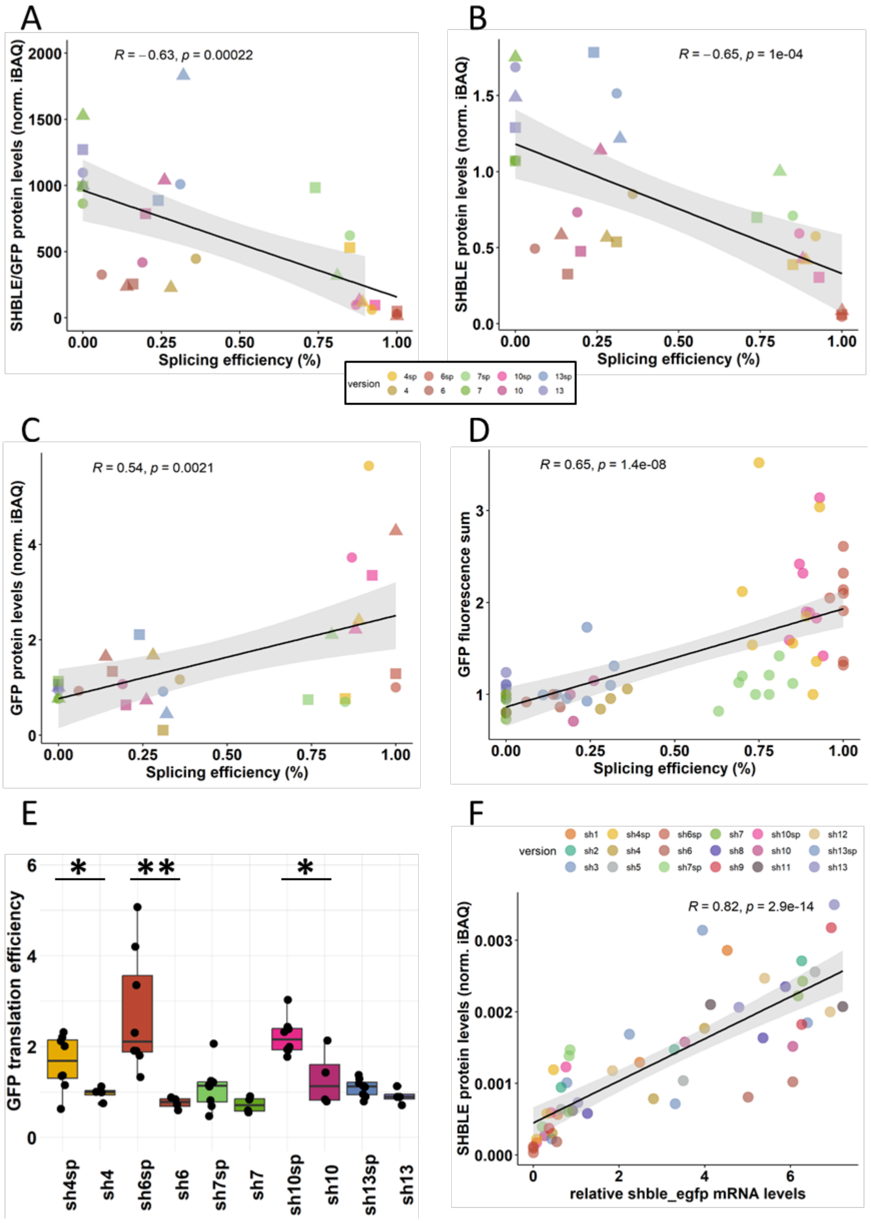


**Figure S9: Splicing effect on SHBLE and GFP translation efficiency. (panel A)** Pearson's linear regression (black line) and 95% confidence interval of the fit (grey) between splicing efficiency and SHBLEoverGFP protein levels expressed in the ten *shble* conditions from three biological replicates. (**panel B**) Pearson's linear regression (black line) and 95% confidence interval of the fit (grey) between splicing efficiency and SHBLE protein levels expressed in the ten *shble* conditions from three biological replicates. **(panel C and D)** Pearson's linear regression (black line) and 95% confidence interval of the fit (grey) between the splicing efficiency and label free proteomic GFP protein levels or cytometry GFP sums of fluorescence expressed in the ten *shble* conditions from three biological replicates. **(panel E)** Box-and-whiskers plot of the GFP translation efficiency calculated from cytometry based sum of fluorescence from splice-able and splice-ablated pairs of construct from four to eight biological replicates. Comparison of paired splice-able and splice-ablated construct revealed statistical differences between pairs of version #4, #6 and #10 after Wilcoxon rank sum test ($\alpha$=0.05, * $p$<0.05, ** $p$<0.01). (**panel F**) Pearson's linear regression (black line) and 95% confidence interval of the fit (grey) between the relative mRNA levels and SHBLE protein levels expressed in the eighteen conditions from three biological replicates. For panels A, B and C, values from a same biological replicate are represented by triangle, rectangle or circle shapes.

### Supplementary Experimental procedures

**Qualitative analyse of heterologous mRNA.** Primers were identical for all constructs, as they targeted transcript regions present in all of them, located in the (FigS1B). Thermal profile consisted of an initial denaturation step of 5min 95 °C, either 25 or 40 cycles of denaturation 30s 95 °C, hybridization 30s 55 °C and elongation 1min 72 °C, with a final elongation step of 5min 72 °C. For detection and identification of spliced isoforms, forty PCR cycles were used (Fig S2A) to maximize amplification of minor heterologous mRNA isoforms and to allow for band extraction for downstream Sanger sequencing. PCR products were analysed on a 1.5% (w/v) agarose gel prepared with 1X TBE (Tris 90mM pH8.3, boric acid 90mM, EDTA 2mM) buffer ran for 45 min at 100V on a 20cm long system, alongside the Generuler 1kb plus da ladder (Fisher Scientific). Revelation was carried on an ebox CX5 UV imager (Vilber). Then, each band was extracted on a UV table and amplicons were purified using the NucleoSpin gel and PCR clean-up kit (Macherey-Nagel) following manufacturer's instructions. Sanger sequencing was carried out by GenoScreen (France) in forward and reverse using 5'UTR-10F and GFP-7R primers, respectively. SANGER profiles were analysed using snapgene viewer v7.1.1 (Fig S2B). Finally, vector and transcript map were constructed on snapgene viewer v7.1.1 using Genscript plasmid supplier information and SANGER data (Fig S1 A and B). For mRNA isoforms quantification 25 PCR cycles were used to remain in the exponential amplification phase for both low and high expressed mRNA isoforms. PCR products were then analysed on a Bioanalyzer (see Experimental procedures).

**Quantitative analyse of heterologous mRNA relative levels.** RTqPCR were performed on 4 µL of 1:20 diluted cDNA, in a reaction volume of 20 µL and analysed on a AriaMx real-time PCR system (Agilent) with a thermal profile consisting of 5 min 95°C followed by 40 cycles of 15sec 95°C and 45sec at 60°C. *B-tubulin* mRNA levels were measured by RT-qPCR and used as internal normalization standards. The $2^{-\Delta\Delta Ct}$ method was used for relative mRNA levels quantification using the human *B-tubulin* gene as internal normalisation control. The same reference sample was added to every RTqPCR plate and was used as reference sample for relative quantification across replicates. All primer and target sequences used for this study tried to be in compliance with MIQE guidelines (Bustin et al. 2009). Primer specificity and efficiency were validated by means of single peak melting curves and single band agarose gel of RT-qPCR amplicons, and serial dilution amplification, respectively.

**Table S2: Primer sequences used**

| primer | sequence | target (RT-qPCR) | (RT-PCR, sanger) |
|---|---|---|---|
| 5'UTR-10F | GAGAACCCACTGCTTACTGG | | Vector insert |
| GFP-7R | GCTTGCCGGTGGTGCAGATG | GFP | |
| GFP-7F | GACGTAAACGGCCACAAGTT | | |
| tubulin-F | TCCTCCACTGGTACACAGGC | B-tubulin | |
| tubulin-R | CTCCTCTTCGGCCTCCTCAC | | |
